# Antibodies to protozoan variable surface antigens induce antigenic variation

**DOI:** 10.1101/2022.06.21.497077

**Authors:** Albano H. Tenaglia, Lucas A. Luján, Diego N. Ríos, Victor Midlej, Paula A. Iribarren, Cecilia R. Molina, M. Agustina Berazategui, Alessandro Torri, Alicia Saura, Damián O. Peralta, Macarena Rodríguez-Walker, Elmer A. Fernández, Juan P. Petiti, Marianela C. Serradell, Pablo R. Gargantini, Vanina E. Alvarez, Wanderley de Souza, Hugo D. Luján

## Abstract

The genomes of most protozoa encode families of variant surface antigens, whose mutually exclusive changes in expression allow parasitic microorganisms to evade the host immune response^1,2^. It is widely assumed that antigenic variation in protozoan parasites is accomplished by the spontaneous appearance within the population of cells expressing antigenic variants that escape antibody-mediated cytotoxicity^1,2^. Here we show, both *in vitro* and in animal infections, that antibodies to Variant-specific Surface Proteins (VSPs) of the intestinal parasite *Giardia lamblia* are not cytotoxic, inducing instead VSP clustering into liquid-ordered phase membrane microdomains that trigger a massive release of microvesicles carrying the original VSP and switch in expression to different VSPs by a calcium-dependent mechanism. Surface microvesiculization and antigenic switching are also stimulated when *Trypanosoma brucei* and *Tetrahymena thermophila* are confronted to antibodies directed to their GPI-anchored variable surface glycoproteins. This novel mechanism of surface antigen clearance throughout its release into microvesicles coupled to the stochastic induction of new phenotypic variants not only changes the current paradigm of spontaneous antigenic switching but also provides a new framework for understanding the course of protozoan infections as a host/parasite adaptive process.

## Main

Antigenic variation (AV) is a mechanism developed by parasitic microorganisms to change the expression of their highly immunogenic surface molecules^1^. AV was discovered more than a century ago^2^ and is considered an intrinsic process responsible for persistent and recurrent infections caused by unicellular microorganisms^1–3^.

*Giardia lamblia* colonizes the lumen of the upper small intestine of vertebrates^4^. Its life cycle includes the environmentally resistant cysts and the proliferating trophozoites, which attach to the gut surface and cause the clinical manifestations of giardiasis^4^. Unlike other protozoan parasites, AV in *Giardia* was discovered as a phenomenon occurring *in vitro^5^* (i.e., in the absence of any immune pressure) before being found to occur during animal infections^6^, suggesting that spontaneous antigenic switching is an inherent characteristic of this parasite.

AV in *Giardia* involves VSPs, which cover the entire trophozoite surface and are the main antigens recognized by their hosts^4–7^. VSPs are integral membrane proteins with antigenically variable ectodomains containing multiple CXXC motifs and a highly conserved C-terminal region, comprising a transmembrane domain (TMD) and a short cytoplasmic tail (CT)^7–9^. The parasite genome encodes a repertoire of 136 VSPs^9^, but only one is expressed on the surface of individual trophozoites at any time^8^. Expression of a unique VSP is regulated post-transcriptionally by an RNAi-like mechanism and maintained epigenetically^10^. However, why and how trophozoites change one VSP for another remains unknown.

As a major host response, the effect of antibodies against surface antigens has basically been studied in terms of their fate after binding to the cell surface. Many protozoa cope with surface-adhered antibodies by either segregating them into “caps” or transferring the antigen/antibody complex to a “flagellar pocket” for endocytosis and lysosomal degradation, thereby avoiding protozoan agglutination and opsonization^11–14^. If antibody concentrations exceed the capability to remove them, parasites might die by either complement-dependent or independent antibody-mediated cytotoxicity^11–16^. Similarly, antibodies against *Giardia* VSPs have extensively been reported to be cytotoxic^6,15–19^.

AV takes place at very low rates in *Giardia* cultures^4,16^ and does not seem to occur in antibody-deficient hosts^4,8^. During the course of infections in immunocompetent individuals, however, if antibody-mediated killing is as efficient as it was found *in vitro*, the number of switchers should be too low to sustain chronic infections. Therefore, we hypothesized that the relationship between protozoan parasites and the host immune response is more complex than previously speculated and that antibodies play a major role during AV.

### Antibodies to VSPs stimulate switching

*Giardia* isolates are arranged into genetic groups or assemblages, each having a particular VSP repertoire ^7–9^. When *Giardia* trophozoites are exposed to anti-VSP monoclonal antibodies (mAbs) or serum collected from infected animals, parasites agglutinate into large aggregates that have been considered clusters of dead cells^6–8,17–19^.

To better explore the effect of antibodies to VSPs, specific mAbs against VSPs from trophozoites belonging to assemblages A1 (isolate WB/C6) and B (isolate GS/M-83) (**Extended Data Table 1**) were used to select by limiting dilution *Giardia* clones expressing either VSP417, VSP1267 or VSPH7. Once cloned (**Extended Data Fig. 1a**), trophozoites were confronted to different concentrations of their specific anti-VSP antibody. Results showed that at high concentrations (100 μM), purified mAbs produced the rapid detachment of the parasites from the culture tubes and their clumping into aggregates containing live trophozoites, which remained grouped but motile for days (**Extended Data Fig. 1b** and **Supplementary Movie 1**). When the same clones were confronted for 72 h to lower concentrations of either their anti-VSP mAb or to an unrelated mAb^20^ (both at 50 nM), anti-VSP antibodies only produced a transient parasite detachment and agglutination (**Extended Data Fig. 1c**), although neither their viability (**Fig. 1a** and **Extended Data Fig. 1d**) nor proliferation (**Fig. 1b**) was affected. However, almost no trophozoite in the anti-VSP antibody treated population maintained the expression of the original VSP, unlike the controls (**Fig. 1c**), indicating that low concentrations of anti-VSP antibodies may induce antigenic switching. Moreover, in clones treated with varying antibody concentrations and for different periods, the percentage of switchers incremented with both increasing induction time and antibody concentration (**Fig. 1e**). Since experiments were performed in clones expressing different VSPs and using a variety of mAbs, the observed effects were not influenced by either the isotype of the anti-VSP mAb or their potential differences in affinity (**Extended Data Table 1**).

**Figure 1.**
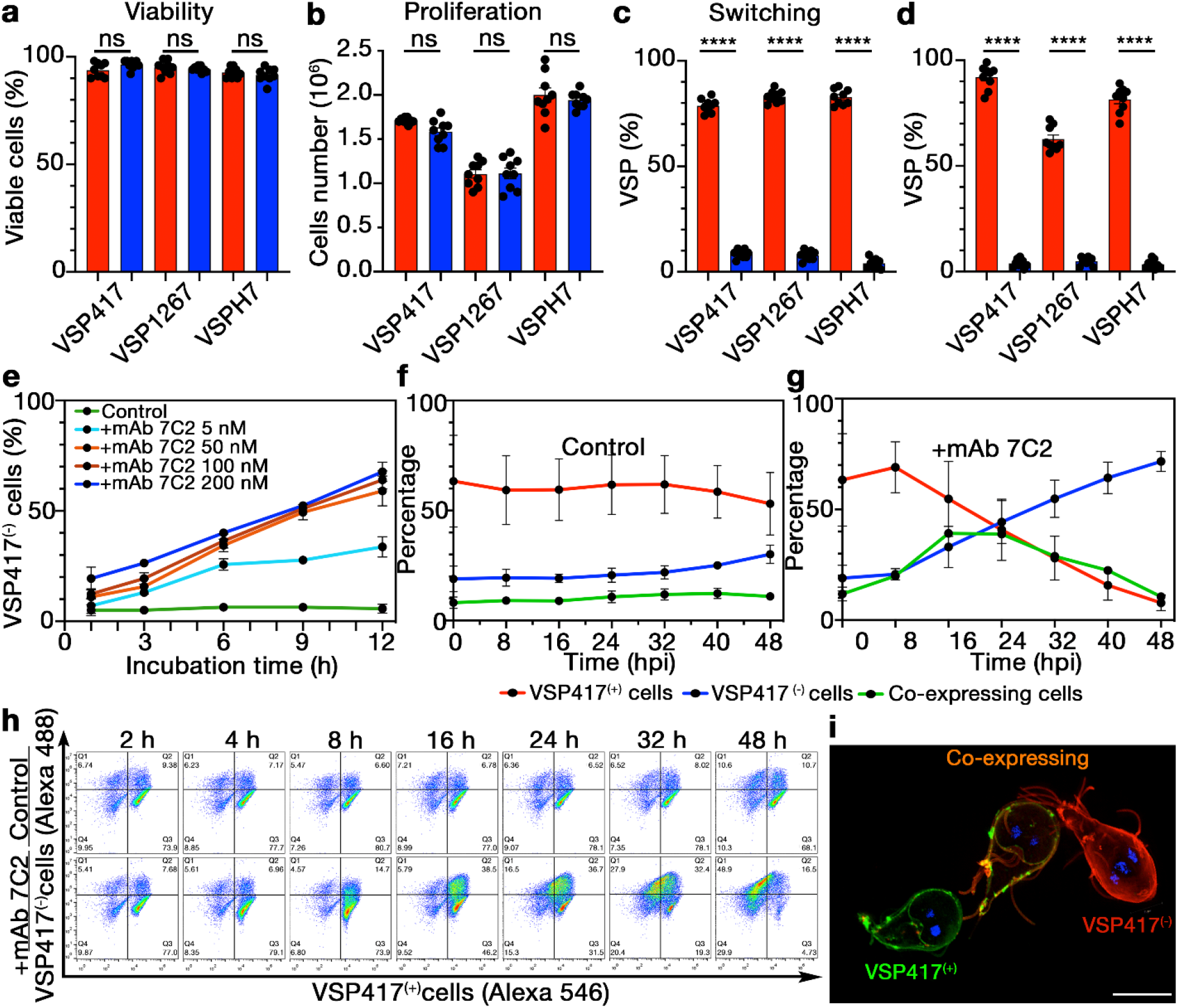
Antibodies to *Giardia* VSPs stimulate antigenic variation. **a-c,** Different VSP clonal populations were grown in the presence of 50 nM of either anti-CWP1 mAb (red columns, controls) or their cognate anti-VSP mAb (blue columns) for 72 h; then, the percentage of live cells (**a**), the total number of cells (**b**) and the percentage of the original VSP in each population (**c**) were determined. **d**, Percentage of cells expressing particular VSPs after *Giardia* cloning in the presence of their cognate anti-VSP mAb (blue) or the control (red). **e**, Percentages of VSP417^(-)^ cells after discrete incubation times with mAb 7C2 at varying concentrations. **f** and **g**, Percentages of VSP417^(−)^, VSP417^(+)^ and co-expressing cells treated with mAb 7C2 (50 nM) for 48 h. **h**, Representative cytograms showing the anti-clockwise distribution shift from VSP417^(+)^ to VSP417^(−)^ cells when cells were stimulated with mAb 7C2 (+ mAb 7C2) at different hpi. Incubation with mAb 8F12 (Control) shows no redistribution. **i**, Super-resolution structured illumination microscopy showing three trophozoites at different stages of antigenic switching after 24 h of mAb stimulation. Scale bar, 5 μm. Values represent mean ± s.e.m. of three independent experiments performed in triplicate. \**p*<0.05; \*\**p*<0.01; \*\*\**p*<0.001; \*\*\*\**p*<0.0001; ns, not significant.

To determine whether the observed AV was caused by induction of switching or by antibody-mediated negative selection, two different approaches were used. First, trophozoites expressing a given VSP were re-cloned in culture medium already containing either the mAbs directed to their specific VSP or the control antibody (50 nM). After 5 days, no differences in the number of growing clones were found (**Extended Data Fig. 2**). Since clones originated from a single trophozoite in contact with their anti-VSP mAb, the absence of antibody-induced killing was confirmed. In the controls, most clones maintained their VSPs, whereas in antibody-treated cells the number of trophozoites expressing the original VSP was particularly low (**Fig. 1d** and **Extended Data Fig. 2**). Moreover, when the expression of different VSPs was examined by immunofluorescence assays (IFAs), all these newly expressed VSPs were present in small percentages, accounting for the absence of any order in the expression of the novel VSPs (**Extended Data Table 2**). Second, the differences in switching rates between clonal populations expressing VSP417 treated or not for 72 h with 50 nM of the anti-VSP417 mAb were evaluated using an adapted Luria-Delbrück fluctuation test^21,22^. At these antibody concentration and incubation time, results showed that anti-VSP antibodies induced a ~72-fold increase in the switching rate as compared to the controls (**Extended Data Table 3**). Besides, the fluctuation^21^ (variance/mean value) was >100 higher in untreated cells than in clonal parasites treated with its corresponding anti-VSP mAb, strongly supporting that AV is induced by antibodies.

To delve into the dynamics of VSP replacement, trophozoites expressing VSP417 were grown in the presence of mAb 7C2 or of the unrelated mAb (50 nM). At different time points, the number of trophozoites labelled with mAb 7C2 and a polyclonal antibody against VSPs other than VSP417 (pAb VSP417^(-)^) was determined by IFAs and flow cytometry. Results showed a decrease of the number of cells expressing the original VSP over time, with the concomitant increase of those simultaneously expressing the original and the novel VSPs, and finally cells expressing only the new VSPs (**Fig. 1f, g**). Representative flow cytometry plots show that control cells exhibited no changes over a 72-h period; in contrast, mAb 7C2-treated trophozoites showed a conversion path from VSP417^(+)^ to VSP417^(-)^ cells throughout the treatment (**Fig. 1h**). This exchange was better observed in three different VSP417-expressing *Giardia* trophozoites incubated for 24 h with mAb 7C2, in which one is still expressing the original VSP, another has already switched to the expression of a new VSP and the third parasite is still in process of switching (**Fig. 1i)**. This period of VSP colocalization clearly shows that there is no negative selection; instead, there is a true exchange of the VSP on the trophozoite membrane.

### Antibody-mediated VSP signalling

If antibodies against the different ectodomains of VSPs stimulated antigenic variation, VSPs might signal throughout their highly conserved C-terminal region, which comprises the TMD and a 5-amino acid CT (**Fig. 2a**).

**Figure 2.**
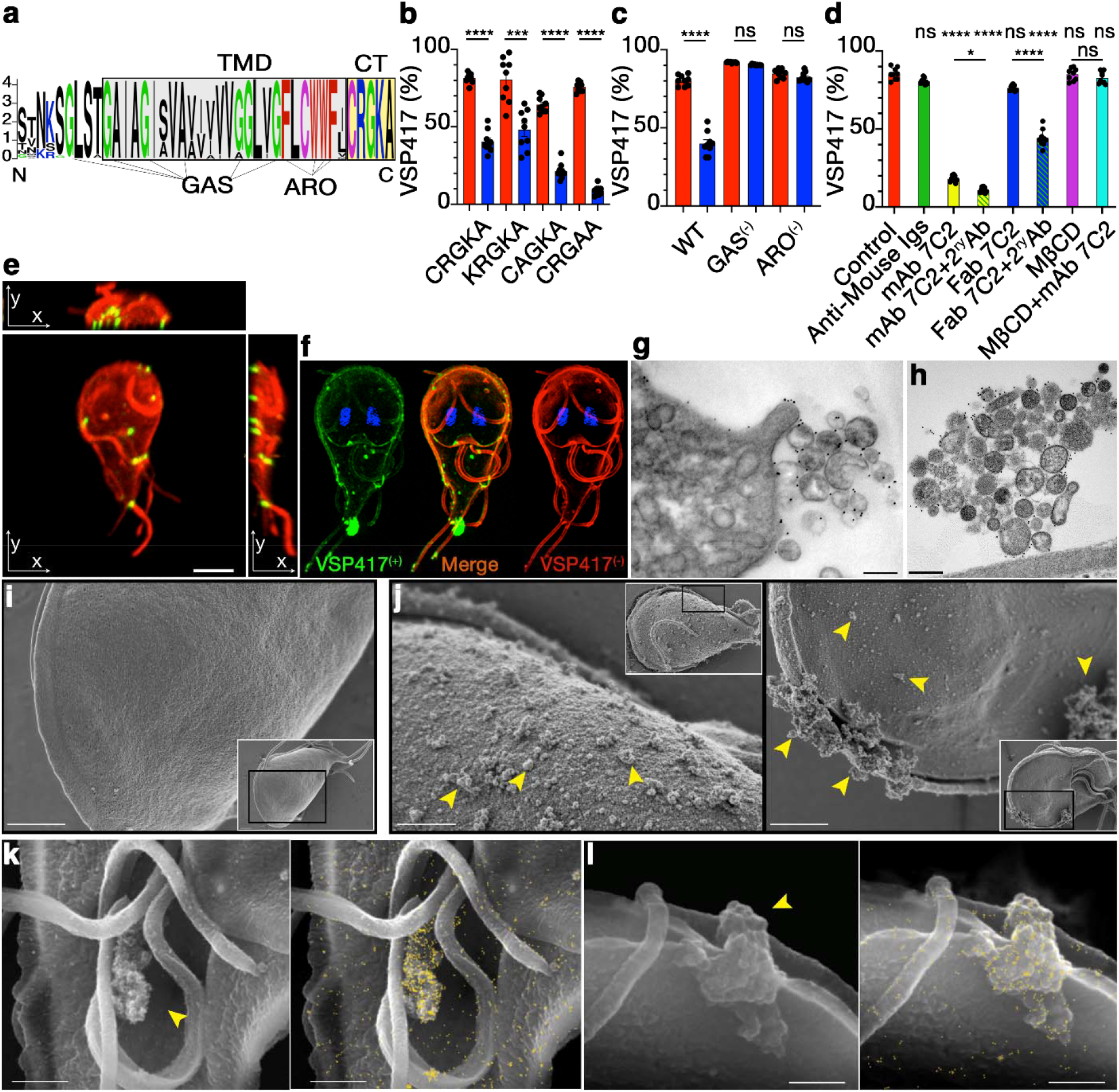
VSP signals through clustering by antibodies. **a**, Sequence logo of the C-terminus of VSPs. TMD and CT are boxed in grey and yellow, respectively. G-xxx-G and small-xxx-small motifs (GAS) and aromatic residues (ARO) are indicated. **b** and **c,** VSP417 trophozoites coexpressing variants of the CT (b) and the TMD (c) of VSPH7 showing the percentage of VSP417^(+)^ cells after 72 h in the presence of either mAb 7C2 (blue) or the control (red). Values were normalized to percentage of cells coexpressing both VSPs. **d**, Percentage of VSP417-expressing cells after 72 h in the presence of mAb 7C2 (50 nM), Fab (100 nM) and Fab in combination with a secondary antibody (2^ry^ Ab), and in the presence or absence of MßCD (10 mM). **e**, VSP417-expressing trophozoite after 1 h of treatment with mAb 7C2 (red); then fixed and incubated with mAb 7C2 (green). Maximum intensity projections on the three orthogonal axes from images taken in Z-stack show microdomains of VSP417 bound to the antibody (yellow) and free VSP417 (red). Scale bar 5 μm. **f**, SR-SIM showing colocalization (yellow) of the former VSP (VSP417, green) and the newly expressed VSP (VSP417^(-)^, red) after 24 hpi with mAb 7C2. **g** and **h**, TEM with gold labelling of a VSPH7 trophozoite incubated for 30 m with mAb G10/4 (g) and of a VSP417 trophozoite incubated for 30 min with mAb 7C2 (h). Scale bars, 500 nm. **i** and **j**, SEM images of a control cell (i) and mAb 7C2-treated trophozoites for 1 h (j). Insets are the cells from which a region (black box) was amplified. Arrowhead shows groups of microvesicles. **k** and **l**, SEM of VSPH7 trophozoites incubated with gold-labelled mAb G10/4 for 30 min showing images obtained by secondary (left) and backscattered electrons (right). Arrows indicate groups of microvesicles containing VSPH7/mAb complexes (yellow dots). Scale bar 1 μm. Values represent mean ± s.e.m. of three independent experiments performed in triplicate. \**p*<0.05; \*\**p*<0.01; \*\*\**p*<0.001. \*\*\*\**p*<0.0001; ns, not significant.

Previous reports suggested the involvement of post-translational modifications of the CRGKA tail of VSPs, such as Cys-palmitoylation and Arg-citrullination in antibody-mediated cytotoxicity, and Lys-ubiquitination in VSP recycling^4,18,19^. Therefore, the influence of the CT of VSPs during AV was evaluated in trophozoites simultaneously expressing two VSPs, which share their conserved C-terminal region but differ in their ectodomains. Trophozoites of the isolate WB endogenously expressing VSP417 were transfected to constitutively express the VSPH7 of isolate GS/M-83 (with or without mutations in its CT) to function as “sensors” of the anti-VSP antibody whereas VSP417 may serve as “reporter” of antigenic switching (**Extended Data Fig. 3a**). Trophozoites co-expressing VSP417 and VSPH7 with and without CT variants (**Extended Data Fig. 3b, c**) were then grown in the presence of mAb 7C2 to induce switching of VSP417 and, separately, in the presence of mAb G10/4 to induce switching of VSP417 through binding to VSPH7. After 72 h, mAb 7C2 promoted switching of VSP417 as expected, but mAb G10/4 also induced switching of the reporter VSP417 in trophozoites co-expressing the VSPH7 wild type CT CRGKA as well as those having the CT variants KRGKA, CAGKA and CRGAA (**Fig. 2b**). These results indicate that, despite its conservation, the short CT of VSPs is not involved in signalling.

Consequently, the highly homologous TMD of VSPs was investigated. A high degree of identity was found between TMD of VSPs and those of surface antigens of other protists (e.g., *Trichomonas* spp., *Spironucleus* spp. and *Leishmania* spp.) and that of the mammalian carcinoembryonic antigen-related cell adhesion molecule 1 (CEACAM1) (**Extended Data Fig. 3b**). The similarity included length and presence of G-xxx-G and small(L/I/V)-xxx-small(L/I/V) motifs (GAS) and a domain rich in aromatic residues (ARO), all known to facilitate the formation of dimers and high-order oligomers^23–25^. Therefore, these motifs were disrupted by substituting the small amino acids of the TMD of VSPH7 for the larger amino acid Methionine and, separately, the aromatic residues of were exchanged for Valines (**Extended Data Fig. 3b)**. Then, the effects of mAb G10/4 in parasites co-expressing VSP417 was analysed. Despite being localized at the cell membrane (**Extended Data Fig. 3c**), TMD variants lacking either the GAS motifs or the ARO domain notably diminished VSP417 switching induced by the anti-VSPH7 antibody (**Fig. 2c**). Since these mutations changed the capability of TMD oligomerization (**Extended Data Fig. 3d**), results indicated that the TMDs of VSPs play an important role in transducing the stimulus for AV.

In addition, since antibodies might facilitate VSP/VSP interactions, mAb 7C2 was fragmented to its corresponding Fabs and their relative ability to induce VSP417 switching was compared. Unlike the intact mAb (50 nM), Fab fragments (100 nM) failed to induce antigenic switching, but their efficiency was restored when Fabs were clustered with secondary antibodies (**Fig. 2d**), suggesting that VSP TMD oligomerization and VSP clustering are crucial for switching.

TMD length and sequence are key determinants of liquid-ordered phase (l_o_) membrane microdomains association^26^ and, indeed, CEACAM1 and oligomerized plasma membrane proteins are usually present on lipid raft-like structures^23–26^, as also are the GPI-anchored variable surface antigens of other protozoa^1,27^. In *Giardia*, the characteristics of VSP TMD suggested that they would also translocate into l_o_-phase, lipid raft-like membrane domains, which were previously described in this parasite^28^. In fact, when trophozoites were cultured in the presence of 10 mM of methyl-ß-cyclodextrin (MßCD, a cholesterol-rich microdomain disrupting agent), VSP switching induced by anti-VSP mAbs was abolished (**Fig. 2d**). Hence, Triton X-100 lysates of trophozoites expressing VSP417 pre-treated or not with 50 nM of mAb 7C2 for 1 h were subjected to sucrose density gradient centrifugation to allow the isolation of low-density detergent resistant membranes (DRM), which have properties of lipid rafts^29^. VSP417 was mainly detected in detergent soluble membranes (DSM) in both treated and untreated trophozoites, but a fraction of the VSP417/mAb 7C2 complexes redistributed into DRM upon antibody binding. Notably, pre-treatment of trophozoites with 10 mM of MβCD before DRM isolation avoided antibody-induced VSP417 redistribution into DRM (**Extended Data Fig. 4**). These results are consistent with results of IFAs, which showed that at 1 h of incubation with the anti-VSP antibody most of the VSP were still present all over the plasma membrane of the trophozoites, whereas the VSP/antibody complex localized into punctate domains (**Fig. 2e**). Therefore, the TMD of the VSPs may direct these proteins into cholesterol-rich microdomains upon antibody binding. Since l_o_-phase domains are dynamic structures, it is likely that VSP-antibody interactions stabilize VSPs into larger platforms capable of recruiting cytoplasmic molecules that activate signal transduction mechanisms^38^.

### VSPs are removed into microvesicles

To track the fate of the VSP bound to its specific antibody, *Giardia* clones expressing different VSPs were treated with their corresponding mAbs and analysed by structured illumination microscopy (SIM) 24 h post-induction (hpi). At the plasma membrane, the cognate VSP formed clusters and the newly expressed VSP showed a more homogeneous distribution (**Fig. 2f**). These microdomains were then studied at the ultrastructural level by combining transmission (TEM) and scanning (SEM) electron microscopy with or without immunogold-labelling. In **Fig. 2g** and **h**, immunogold TEM micrographs of a *Giardia* trophozoite at 30 mpi showed groups of microvesicles (MVs) on their surface. By SEM, the cell surface changed from smooth in untreated trophozoites (**Fig. 2i**) to rough, full of MVs in antibody-treated cells (**Fig. 2j**). SEM with gold labelling showed that these MVs clumps were enriched in the target VSP (**Fig. 2k, l**). In contrast to the current assumption that the former surface antigen disappear by its continuous dilution in the plasma membrane after a new antigen is being expressed, these results indicate that after antibody binding, VSPs are eliminated from the cell surface into extracellular MVs. In this process, called “<u>A</u>ntigen <u>R</u>emoval <u>C</u>oupled to <u>S</u>witching (ARCS)”, both surface antigen clearance and antigenic variation are simultaneously triggered by surface antigen clustering into lipid raft-like microdomains.

To test if a similar process occurs *in vivo*, gerbils (*Meriones unguiculatus*) were infected with VSP417-expressing trophozoites and intestinal parasites were collected and analysed by SEM and flow cytometry at different days post-infection (dpi). At the onset of the humoral immune response (**Fig. 3a**), the same protrusions and MVs that were seen upon antibody treatment *in vitro* were observed in trophozoites collected from the small intestine (**Fig. 3b, c**). Moreover, conversion from VSP417^(+)^ to VSP417^(−)^ cells was observed between 10 and 12 dpi, with a peak of original/new VSP co-labelling at 11 dpi (**Fig. 3d**). As observed *in vitro*, switching to different VSPs within the intestine was random (**Fig. 3e**).

**Figure 3.**
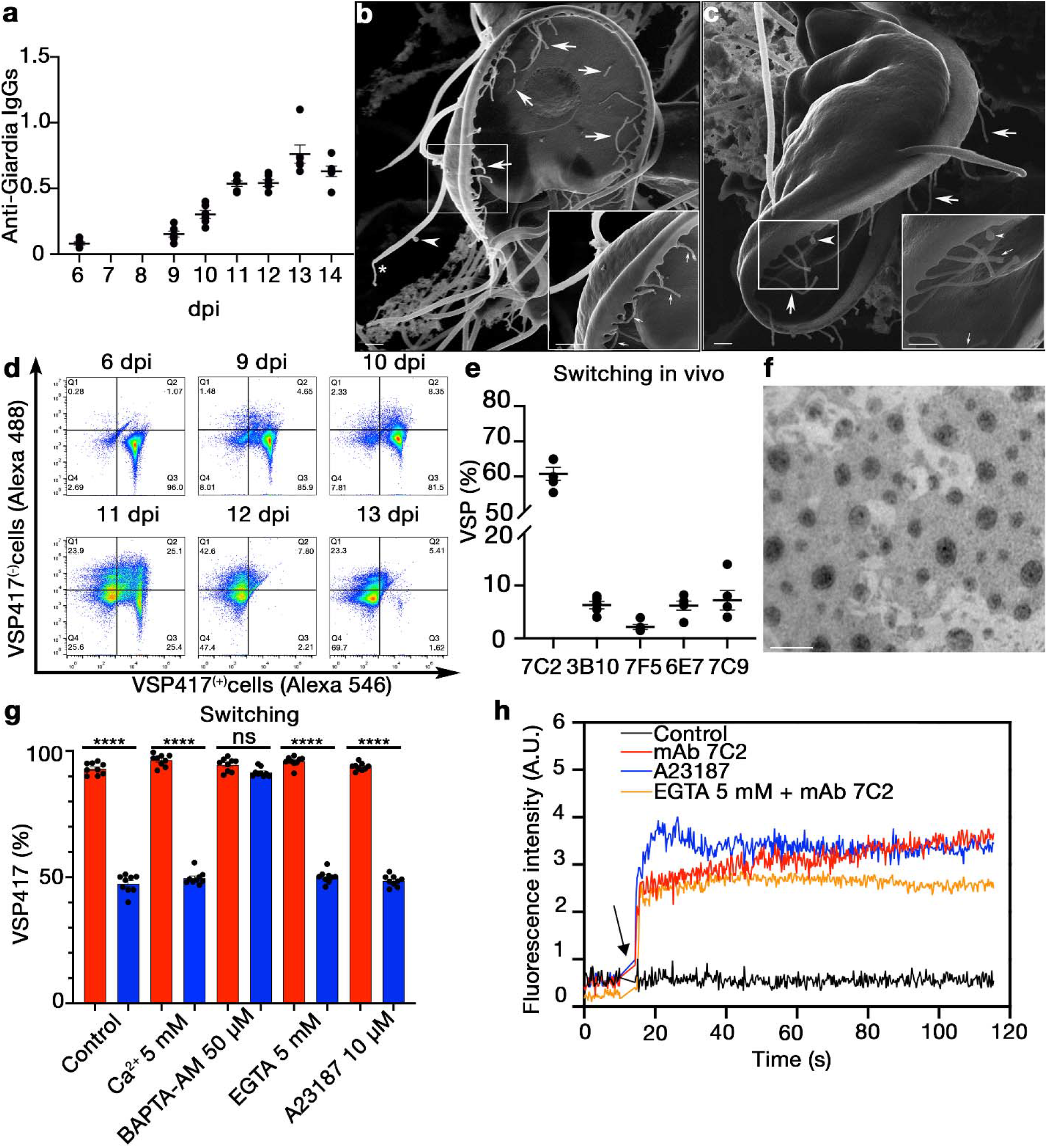
VSPs are released into microvesicles. **a**, Total antibodies against *Giardia* VSP417-expressing trophozoites detected by ELISA in sera from infected gerbils at different dpi. Values represent relative optical density. **b** and **c**, Extreme Resolution HI-SEM of VSP417 trophozoites recovered from the small intestine at 12 dpi. Extensive membrane projections emerging from the cell flange and the ventral disc are shown (arrows). A vesicle budding from a flagellum (arrowhead) and membrane projections at the end of a flagellum (asterisks) are observed. Scale bar 1 μm. **d**, Representative cytograms showing the distribution shift from VSP417^(+)^ to VSP417^(-)^ populations during animal infections with a VSP417 clone. **e**, Percentage of intestinal trophozoites recognized by different anti-VSP mAbs at 11 dpi. **f**, Negative staining of purified microvesicles obtained trophozoites of clone VSP417 incubated with mAb 7C2 for 4 h. Scale bar, 200 nm. **g**, Effects of different concentrations of extracellular calcium, EGTA, BAPTA-AM and the Ca^2+^ ionophore A23187 on antigenic switching of VSP417-expressing trophozoites grown in the presence of a control mAb (red) or of mAb 7C2 (blue) for 72 h. **h**, The effects of anti-VSP mAb 7C2, a control antibody and the calcium ionophore A23187 on the intracellular levels of Ca^2+^ in VSP417-expressing cells are shown. Each line corresponds to the mean value of three independent experiments. Arrow indicates the time at which the stimulus was added. Values in other panels represent mean ± s.e.m. of three independent experiments performed in triplicate. \**p*<0.05; \*\**p*<0.01; \*\*\**p*<0.001; \*\*\*\**p*<0.0001; ns, not significant.

To determine the composition of the MVs, proteomic analysis (**Extended Data Table 4**) of purified MVs collected 4 hpi (**Fig. 3F**) demonstrated that their most enriched molecule is the original VSP. Except for multiple annexins (α-giardins^30^), ALIX/PDC6IP^31^ and components of the ESCRT machinery^32^, most proteins found in the antibody-induced MVs (**Extended Data Table 4**) were similar to *Giardia* MVs artificially generated by incubation of the parasites with 2 mM of extracellular calcium^33^.

To test if antigenic variation can be triggered by the sole induction of MVs, clonal trophozoites were incubated with different extracellular [Ca^2+^] in either absence or presence of anti-VSP antibodies. Neither viability nor proliferation was affected after 72 h of incubation, except in the presence of high concentrations of EGTA (10 mM) (**Extended Data Fig. 5**), but VSP switching was only induced when antibodies directed to their cognate VSP were present (**Fig. 3g**). Conversely, the intracellular Ca^2+^ chelator BAPTA-AM (0.5 μM) suppressed switching in trophozoites treated for 72 h with antibodies against their VSP (**Fig. 3g**), without affecting parasite viability or proliferation (**Extended Data Fig. 5**). To determine if VSP clustering by antibodies promotes an increase in the intracellular [Ca^2+^], trophozoites expressing VSP417 previously loaded with the calcium indicator Fluo 4-AM were subsequently incubated with either mAb 7C2 or a control mAb (50 nM). Results showed a strong increase in the intracellular [Ca^2+^] soon after antibody binding to the VSP (**Fig. 3h**), in the presence or the absence of extracellular Ca^2+^. Remarkably, although the Ca^2+^ ionophore A23187 (10 μM) also increased the intracellular [Ca^2+^] to similar levels (**Fig. 3h**), it did not induce antigenic switching (**Fig. 3g**). This result suggests that an intracellular Ca^2+^ increase is necessary but insufficient to trigger ARCS and that the mechanical clustering of VSP caused by anti-VSP antibodies elicit a more complex process responsible for antigen elimination and switching. Since Ca^2+^ is essential for the function of annexins, ALIX/PDCD6IP and PDCD6/ALG-2 (**Extended Data Table 3**) during plasma membrane repair, virus budding and signal transduction^34–36^, these results suggest that anti-VSP antibodies promote the discharge of Ca^2+^ from intracellular stores to promote ARCS.

### ARCS occurs in other protozoa

The fact that the TMD of VSPs makes these molecules raftophilic highlights the similarity to other protozoan variable surface antigens that are attached to the plasma membranes by GPI anchors^1,27,37^. Consequently, antibodies to *T. brucei* variant surface glycoproteins (VSGs) and *T. thermophila* i-Antigens were generated and used to determine if a process similar to that found in *Giardia* is also occurring in these protozoa.

In *T. brucei*, mouse anti-VSG antibodies generated during experimental infections were not cytotoxic at high dilutions (>1/1,000) and turned the parasite smooth surface into one full of nanotubes and MVs after antibody treatment (**Fig. 4a-f**). Since these morphological changes were similar to those observed in *Giardia* trophozoites exposed to anti-VSP antibodies, the connexion between cell membrane microvesiculation induced by anti-VSG antibodies and antigenic switching was evaluated using two different procedures. First, an experimental method was designed to test the influence of increasing concentration of anti-VSG antibodies on the growth of *T. brucei* and to allow the identification of potential switchers (**Extended Data Fig. 6**). Briefly, three parallel cultures of parasites expressing VSG AnTat1.1 were grown for 72 h in the presence of either preimmune serum or sub-lethal daily doses of the anti-VSG pAb. Then, serial dilutions of each culture were spread over a 96-well plate and parasites were exposed to a lethal dilution of the anti-VSG pAb (1/50) to kill non-switchers. After 6-8 days, wells containing live parasites were subjected to cDNA sequencing for identification of switchers. Results demonstrated that increasing concentrations of specific anti-VSG antibodies induce switching to different VSGs without affecting proliferation of the parasites (**Fig. 4g**). Furthermore, while the two clones that survived in the control expressed the same VSG, clones isolated from the cultures treated with the anti-VSG pAb expressed a diversity of VSG (**Fig 4g**), confirming that they indeed correspond to independent switching events. Second, to determine switching rates in the presence or absence of anti-VSG pAb, an adapted Luria-Delbrück fluctuation test^21^ was performed using either lethal dilution of the anti-VSG antibody (1/50) or a well described *VSG* RNAi strategy^22^ for the selection of switchers. In these experiments, parallel cultures containing a very low number of parasites were expanded for 8-9 generations in the presence of either anti-VSG AnTat1.1 pAb (1/1,000) or the preimmune serum before cultures were spread over 96-well plates for selection of switchers by using a lethal dilution of the anti-VSG pAb (1/50) or, separately, by addition of doxycycline to induce *VSG* AnTat1.1 RNAi. Since *VSG* RNAi results in cell-cycle arrest and subsequent death, only parasites that have switched to a new VSG variant will be able to grow^22^. Using both approaches, the calculated switching rate of the parasites treated with anti-VSG pAb increased 10-25 times as compared to the parasites incubated with the control serum (**Extended Data Table 5a and b**). Furthermore, the variance in the number of clones generated per culture divided by the mean resulted much closer to 1 in the presence of anti-VSG pAb (**Extended Data Table 5a and b**), confirming that VSG switching is induced by the anti-VSG antibodies.

**Figure 4.**
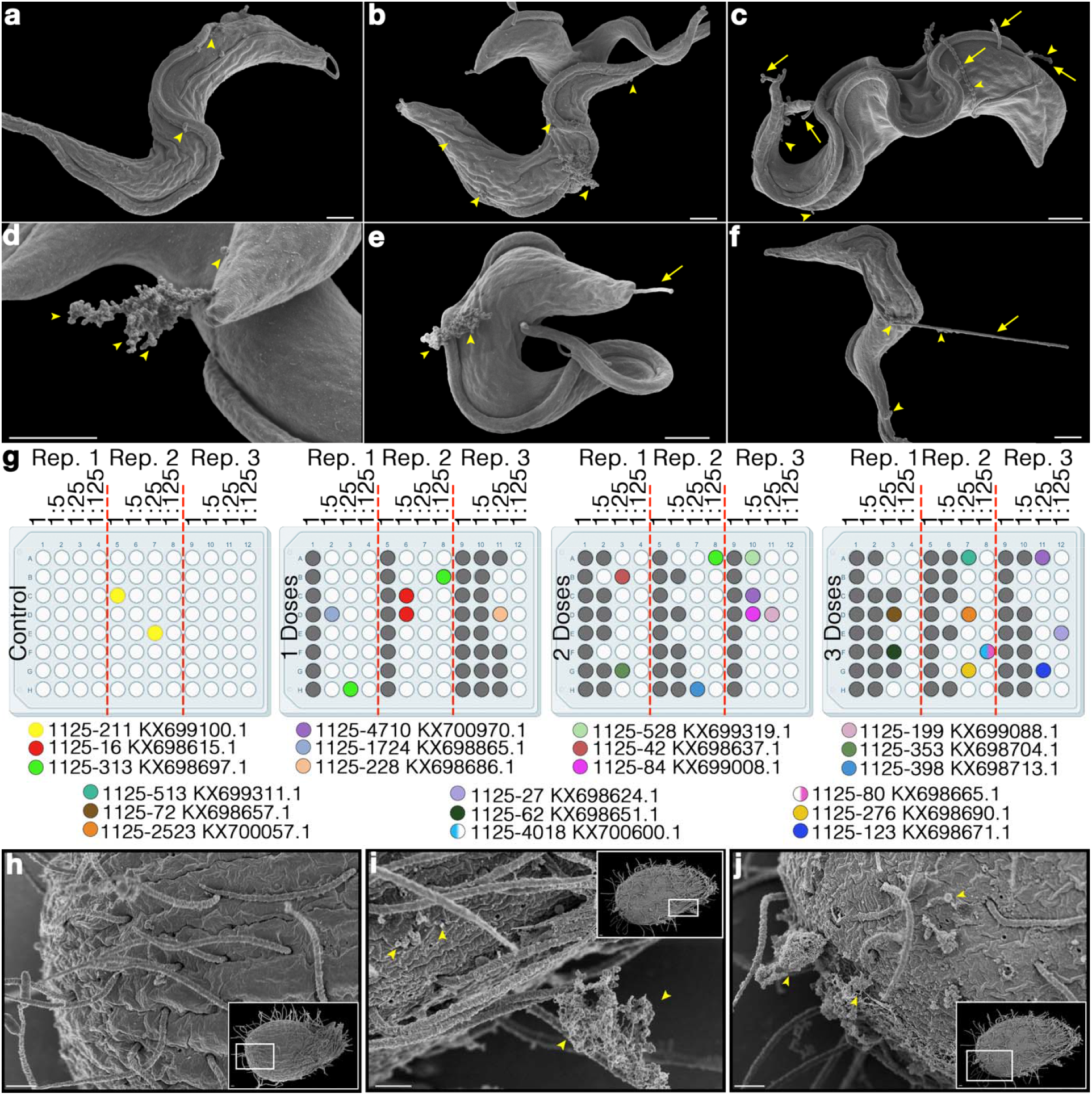
ARCS in other protozoa. **a-f**, HR-SEM micrographs of monomorphic forms *T. brucei* expressing VSG 221a incubated with mouse serum obtained 15 dpi at 1/1,000 dilution (a, 5 mpi, b-f, 4 hpi). Extensive membrane projections emerging from the cell surface and large amounts of microvesicles can be observed (arrowheads). Scale bars 1 μm. **g**, Induction of switching by three daily additions of anti-VSG antibodies in *T. brucei* clone EATRO1125 expressing VSG AnTat1.1, following the protocol depicted in Extended Data Fig. 6. After 72 h, non-switchers were killed with a lethal antibody dilution (1/50) and switchers were selected at random in each plate for their identification (the first number or letter of the ID code indicates the plate: control; 1 dose; 2 doses; 3 doses; the following letter and number indicate the position within the plate) along with the identified VSG and a unique colour code for each VSG. Wells in which the VSG was not identified are in grey. **h**-**j**, HR-SEM micrographs of *T. thermophila* cells untreated (**h**) or treated with an antiserum directed to their variable surface antigens after 4 hpi (**i, j**). Insets correspond to the whole cells and the amplified image is denoted by black boxes. A massive release of microvesicles still attached to the cell surface can be observed in treated cells, which is not seen in the controls. Scale bars 2 μm.

Similar experiments were performed in clones of the free-living protozoan *T. thermophila*. High dilutions (>1/1,000) of antibodies to their surface antigens generated a temporary agglutination of the cells with the concomitant disappearance of the recognized antigen, which reappeared on their surface several days after the antibody was removed from the culture (**Extended Data Fig. 7**). As in *Giardia* and *T. brucei*, SEM analysis showed that the surface of this ciliate appeared decorated with clumps of MVs soon after antibody treatment (**Fig. 4h-j**), clearly denoting that ARCS may be a widespread process in protozoa.

## Discussion

Antigenic variation seems to occur sporadically in protozoan cultures, which was interpreted as a spontaneous process that allows switching of surface antigens after a given number of generations^1^. Unlike previously assumed^6,15–19^, antibodies against variant surface antigens are not cytotoxic at low concentrations; rather, they strongly stimulate antigenic switching. This is the first description of a host-induced tuning of AV, with important consequences for understanding the course of protozoan infections and the propensity for multiple reinfections.

Our findings show that variable surface antigens acquire the role of sensors of the humoral immune response, promoting antibody-directed phase separation of antigens into l_o_-phase microdomains. The mechanical stress of the plasma membrane caused by antibody binding to raftophilic variable surface antigens then triggers the immediate clearance of the former antigen into extracellular MVs while stimulating antigenic switching. Although the induction of microvesicles by different stress conditions has been observed in many protozoan parasites^38^, their association with AV has never been investigated. Our novel observations that the original surface antigen is removed from the plasma membrane into microvesicles suggest that this process may divert the immune response to these MVs to allow the parasite to complete antigenic exchange before increasing concentrations of antibodies agglutinate the cells and/or exceed their endocytic capacity.

In the intestinal parasite *G. lamblia*, anti-VSP antibodies may cause not only antigenic switching at early stages of antibody production by the host but also parasite detachment from the gut epithelium when antibody concentrations increase during infections. Due to peristalsis, agglutinated trophozoites can thus reach the lower portions of the small intestine, where they differentiate into cysts^4^. Released cysts containing trophozoites already expressing different VSPs are then prone to reinfect the same individual and to expand the range of potential hosts^8,39^.

In contrast to *Giardia*, in which higher antibody concentrations favour parasite clumping and thus cell differentiation, increasing concentrations of anti-VSG antibodies may kill most of the bloodstream forms of *T. brucei* before some switchers can proliferate, shaping infections as multi-waves instead of an ever-growing curve expected for a survival strategy. It is known that VSG switching involves enzymes of the phosphoinositol phosphate pathway^40^ and that antibodies to VSGs also trigger a cytoplasmic increase of calcium levels^41^, suggesting that acidic phospholipids and Ca^2+^ might also be involved during antigenic variation in this parasite. Moreover, antibody-induced switching in *T. brucei* may explain why the *in vivo* dynamics of infections shows a higher VSG diversity than that observed *in vitro*^42^ and how antibodies lose their ability to mediate parasite clearance at unexpectedly early stages of VSG coat replacement^43^.

Since many pathogenic protozoa regulate the expression of a unique variant by post-transcriptional and epigenetic mechanisms^10,44^, antibodies might reset these control systems through an increase of intracellular [Ca^2+^]^45,46^, which may likely occur occasionally in culture. However, regardless of the mechanisms each pathogen uses to switch its surface antigens, one tempting hypothesis is that the antibody-stimulated release of MVs produces stochastic changes in the cytoplasmic levels of specific molecules in individual cells, resetting the epigenetically controlled antigen-expression program. This possibility might explain the stochastic nature of the switching response as well as the need for an increasing antibody exposure to complete antigenic switching of the whole parasite population^47^.

In *Tetrahymena* and other free-living protozoa, antigenic variation was shown to occur in response to environmental changes^27^. Therefore, the existence of ARCS in such divergent microorganisms highlights the evolutionary significance of this process for protozoan adaptation from a free-living to a parasitic lifestyle, or vice versa^48^.

## Supporting information

Extended data with figures

## Acknowledgments

Authors thank Dr. Theodore E. Nash (NIAID, NIH, USA) for providing mAbs G10/4 and 6E7, Sergio R. Oms for technical assistance and animal care, and Dr. Marcela de Freitas Lopes (UFRJ, Brazil) for critical reading of the manuscript.

## Author contributions

AHT, LAL and DNR contributed the *in vivo* and *in vitro* antigenic switching experiments in *Giardia*. LAL and CRM contributed to the generation of mutants, calcium experiments and fluctuation tests. AT contributed pilot *Giardia* viability, proliferation and switching experiments. AS generated anti-VSP mAbs and pAbs against *T. brucei* and *T. thermophila*. DOP performed mAb and antibody fragment purification and quantification. MCS contributed to flow cytometry experiments. MRW and EAF performed statistical analyses, selected sample sizes and reanalysed proteomics results and fluctuation tests. PAI, MAB, AHT, LAL, CRM, PRG and VEA contributed the *T. brucei* experiments; VM and WDS contributed the confocal, scanning and transmission electron microscopy; JCP validated purified MVs. VEA, WDS and HDL analysed the data. HDL conceived the project and designed the experiments. AHT, LAL, DNR, VEA, WDS and HDL wrote the paper. All authors read and made comments on the manuscript.

## Data and materials availability

All data are available in the main text or the supplementary materials. Proprietary antibodies are available upon request through a material transfer agreement (MTA). The mass spectrometry proteomics data have been deposited at the ProteomeXchange Consortium via the PRIDE partner repository with the dataset identifiers PXD031141 and 10.6019/PXD031141 (Reviewer’s account details: Username; Password: QmM5xe8V).

