## Extended data with figures for "Antibodies to protozoan variable surface antigens induce antigenic variation"

**Methods**

**Protozoan cultures**

*Giardia lamblia* assemblage A1 isolate WB (ATCC^®^ 50803) clones and clone GS/M-83 (ATCC^®^ 50581) (assemblage B) were cultured in TYI-S-33 medium, as previously described^1^. Clones expressing different VSPs were obtained by limiting dilution in 96-well culture plates placed in anaerobic chambers (Anaerogen^TM^ Compact, Thermo Scientific^®^ Oxoid^®^, Cat. # AN0010C) at 37°C for 5 days and positive clones were then selected using specific anti-VSP mAb by immunofluorescence assay (IFA) as described^2^. Reactive clones were then expanded in culture medium overnight and tested for homogeneity before use^3^. For in-cell studies, these clones were re-cloned as described above, but in the presence of 50 nM of their specific anti-VSP mAb. *Tetrahymena thermophila* WH-6 (WHI) (ATCC^®^ Cat. # 16539) was maintained and cloned in standard PPYE medium at room temperature^4^ and T*. brucei* was maintained and cloned as previously described^5,6^. Monomorphic forms of *T. brucei* *brucei* Lister 427 clone 221a were used for morphological analysis, whereas pleomorphic forms of clone EATRO1125 expressing VSG AnTat1.1 (VSG 1125.4156, KX700731.1) were used for switching assays^7^. The *VSG* RNAi cell lines are transformants of clone EATRO1125 expressing VSG AnTat1.1. The RNAi construct contains two AnTat1.1 gene-derived segments inserted in opposite orientations targeted into the ribosomal locus by the *Tb*FIX vector^8^. Thus, transcription driven by the inducible rRNA promoter would produce a double stranded RNA molecule in a “stem-loop” configuration. All media contained antibiotic/antimycotic solution at the recommended concentration (Gibco^®^, Cat. # 15240062).

**Animals**

BALB/c mice, Wistar rats and gerbils (6-8 week-old) of both sexes were housed in the vivarium of the CIDIE under specific pathogen-free (SPF) conditions in microisolator cages (Techniplast^TM^), following NIH guidelines for laboratory animals. All procedures followed the protocols approved by the Institutional Committee for Care and Use of Experimental Animals (CICUAL protocols CIDIE.2016-36-15p-2, and CIDIE.2018-36-15p-3).

**Infections with *G. lamblia***

Before infection, six-week-old gerbils were tested for negativity of serum antibodies against *Giardia* antigens by ELISA, as previously reported^1^. Infections were carried out by orogastric administration of 2x10^5^ trophozoites expressing a particular VSP, which were resuspended in 500 µl of PBS. Trophozoites from the intestine were collected at different days post-infection. Gerbils were sacrificed in a CO_2_ chamber and a 14-cm portion of the upper small intestine (measured from its junction to the stomach) was removed, dissected longitudinally using surgical scissors, incubated in 7 ml of PBS on ice for 30 min and vortexed to detach the parasites. The total number of parasites was counted from the intestinal content suspension using a Neubauer chamber. A centrifugation step at 1,000 g for 5 min was performed to isolate parasites from the intestinal content. PBS was discarded and replaced with culture medium at 37°C. The parasites were incubated at 37°C for 40 min to allow attachment to the walls of the culture tube; the medium was then removed along with the remains of intestinal content. Fresh medium was added and the tubes were incubated on ice to detach the parasites for IFA and electron microscopy analyses.

**Production of monoclonal antibodies**

Mouse mAbs to individual VSPs were generated as previously reported^9^. The isotype and light chain composition of these antibodies were determined using a commercial kit (mouse monoclonal antibody isotyping kit, dipstick format; Bio-Rad, Cat. # MMT1). Monoclonal antibody purification was carried out by FPLC using the ÄKTA^TM^ Pure 25 apparatus (GE Healthcare Life Sciences) coupled to a HiTrap^TM^ Protein G HP (GE Healthcare^®^, Cat. # 29-0485-81), following the manufacturer's protocol. After purification, a change from buffer to PBS was performed using the same equipment coupled to the HiPrep^TM^ 26/10 desalting column (GE Healthcare, Cat. # 17-5087-01). Subsequently, protein concentration was measured using the BCA Protein Assay Kit (Pierce^®^, Cat. # 23225). The concentrations, expressed in molarity, are derived from the protein concentration present in the purified antibodies, considering a molecular weight of 150 kDa (for IgG). Purified antibodies were stored in small aliquots at -20°C.

**Production of polyclonal antibodies**

For rat anti-VSP417^(-)^ polyclonal antibody production, 8-week-old Wistar rats were immunized intraperitoneally on days 0, 14, and 28 with 50 µg of total protein extract of trophozoites derived from clones expressing VSP417 cultured during a week in the presence of 1 µM of mAb 7C2 to ensure complete switching to different VSPs. Protein extracts were emulsified in Sigma Adjuvant System (Sigma-Aldrich, Cat. # S6322) before immunization. Rats were boosted intravenously on day 35 with 25 µg of the protein extract. Three days later, rats were euthanized and total blood was withdrawn by cardiac puncture. Polyclonal serum was obtained and tested as non-reactive to VSP417-expressing trophozoites and reactive to most non-VSP417-expressing cells by IFA. VSP417^(-)^ pAb labelling for IFA and flow cytometry were performed on live, freshly collected trophozoites as described below. Polyclonal antisera against *T. thermophila* and *T. brucei* were produced in 6-week-old BALB/c mice. For *Tetrahymena*, animals were immunized as described above with 30 µg of total protein extracts and the polyclonal serum was tested by IFA for surface labelling. For *T. brucei*, mice were infected intraperitoneally with 1.5x10^6^ parasites. After 48 h, mice that presented detectable parasitemia were cured with two consecutive doses, 24 h apart, of 40 mg/kg of Berenil^TM^ (MSD Animal Health^®^, Reg. SAGARPA Q-0273-121). Visualization and culture of the blood of the animals showed no presence of parasites after Berenil^TM^ treatment. Then, on days 15 and 30, blood samples were taken from the retro orbital plexus using heparinized capillaries. From the obtained sera, the presence of reactive antibodies against the surface of the parasites was confirmed by IFA. Monomorphic forms were used for electron microscopy experiments and the pleomorphic forms were used for switching induction experiments.

**Flow cytometry**

*Giardia* trophozoites were analysed by flow cytometry using a BD Accuri C6^TM^ instrument (BD Biosciences^®^). Before acquiring the data, 8-point beads were used for manual quality control of the instrument. Unstained cells and compensation beads (BD Biosciences, CA) were used to set voltages and create single stain negative and positive controls. Compensation was set to account for spectral overlap between the two fluorescent channels. No gating strategy was used but regions were set as quadrants reflecting positive staining for VSP417, positive staining for VSPs other than VSP417, and positive for both dyes. Density plots were displayed indicating the percentage of each population. Data analysis and graph generation was performed using FlowJo^TM^ 7.6 software (TreeStar^®^).

**Generation of Fab fragments**

Production of Fab fragments from mAbs 7C2 and 7F5 (both isotype IgG1) was performed using an antibody Fab preparation kit from Pierce^®^ (Thermo Scientific^®^, Cat. # 44985) according to the manufacturer's instructions. The Fab fragments were purified by FPLC using the ÄKTA^TM^ Pure 25 equipment coupled to a HiScree^TM^ MabSelec^TM^ SuRe^TM^ affinity column (GE Healthcare^®^, Cat. # 28-9269-77), according to the manufacturer's protocol. The integrity of the binding sites of the Fab fragments was verified by IFA on *Giardia* trophozoites expressing the corresponding VSPs. The absence of undigested mAb was verified by SDS-PAGE under reducing and non-reducing conditions. The Fab concentrations expressed in molarity were derived from the protein concentration present in the purified ones, considering a molecular weight of 50 kDa.

**Clonality assessment**

For *Giardia*, IFA was performed as previously reported^1-3^. The percentage of cells expressing particular VSPs was calculated by counting 500 x 3 cells in triplicate experiments or by flow cytometry using an Accuri^TM^ C6 flow cytometer (BD Biosciences^®^). For *T. thermophila***,** cells were labelled alive by incubation with the polyclonal serum at a dilution of 1/1,000 in culture medium. Subsequently, after two washes with PBS, cells were fixed with 4% paraformaldehyde at room temperature for 1 h, washed 3 times with PBS, and incubated with 50 mM NH_4_Cl in PBS for 10 min. Blocking was performed with 3% bovine serum albumin (BSA) solution in PBS for 1 h and then cells were incubated with Alexa Fluor^TM^ 488-labelled goat anti-mouse antibody 1/1,000 in 1% BSA in PBS at room temperature for 1 h. Finally, after three washes, cells were resuspended in ES Glycerol Mounting Medium with DABCO™ (EMS Cat. #17989-50) for IFA. For *T. brucei*, cells were labelled with polyclonal antisera 1/500 in culture medium on ice for 40 min. Subsequently, after two washes with TDB (*Trypanosome* dilution buffer, KCl 5 mM, NaCl 80 mM, MgSO_4_ 1 mM, Na_2_HPO_4_ 20 mM, NaH_2_PO_4_ 2 mM, Glucose 20 mM pH 7.7), parasites were fixed in suspension with paraformaldehyde 4% in PBS at RT for 1 h. The parasites were subsequently adhered to coverslips using a poly-L-lysine solution (Sigma, Cat. # P4707), washed and blocked as described above. Cells were incubated with Alexa Fluor^TM^-546 goat anti-mouse IgG (H+L) (Invitrogen^TM^, Cat. # A11030) or Alexa Fluor^TM^-488 goat anti-mouse IgG (H+L) (Invitrogen^TM^, Cat. # A11003) antibodies at 1/1,000 dilutions. For all parasites, nuclei were stained with DAPI at a concentration of 1 µg/ml in PBS for 10 min. For IFA, images were acquired using a Leica IRBE epifluorescence microscope with a 63X objective (using oil immersion; OA 1.4), equipped with an ORCA ER-II CCD camera (Hamamatsu). Images and videos were acquired using HCI Hamamatsu software. Image analysis and colocalization were done with ImageJ software.

**Cytotoxicity and proliferation assays**

*Giardia* trophozoites (1x10^4^) of clones VSP1267, VSP417 and VSPH7 were incubated with 50 nM of its corresponding mAbs (7F5, 7C2 and G10/4, respectively) during 72 h; then, 100 µl were used to analyse viability by propidium iodide/fluorescein diacetate staining^10^. The other 100 µl were used to count the total cell number in a haemocytometer. Similar procedures were followed with *T. thermophila*. However, to determine the cytotoxic capability of anti-VSGs anti-sera on *T. brucei*, a titration of the sera obtained after 15 dpi was performed. In contrast to *Giardia*, high serum concentrations were lethal to the parasites (dilutions <1/500) and low serum concentrations (dilutions >1/500) transiently agglutinated the parasites that then switched their VSG.

**Populational switching assay**

*Giardia* trophozoites (1x10^4^) of clones expressing VSP417, VSP1267 or VSPH7 were grown under the continuous presence of their cognate anti-VSP mAb (0.25-10 nM) in TYI-S-33 medium. After 1-3 days of culture, the number of trophozoites was quantified in a haemocytometer and VSP characterization was performed by IFA. Double Fab molar concentrations were used to ensure equal amounts of binding sites. For transient-stimulus-assay, 1x10^4^ trophozoites were treated with 50 nM of their corresponding anti-VSP mAb or mAb 8F12, or in 100 nM of Fabs in TYI-S-33 medium for 1 h on ice, washed, and cultured under normal conditions for 3 days before VSP characterization.

**In-cell switching assay**

Fifty trophozoites expressing a particular VSP (VSP417, VSP1267, and VSP2B10) were diluted in 20 ml of TYI-S-33 medium containing 50 nM of the corresponding anti-VSP mAb and distributed in 96-well plates. The same procedure was followed in TYI-S-33 medium with an unrelated mAb as a control (mAb 8F12). Plates were incubated for 5 days; then, the number of clones obtained was counted and the percentage of the original VSP was measured by IFA. For experiments with different incubation times and mAb concentrations, VSP417^(+)^ trophozoites were treated with 0, 5-500 nM of mAb 7C2 in TYI-S-33 medium at 37°C for 1, 3, 6 and 12 h. After incubation, VSP417 expression was determined by IFA.

**Switching rate determination in *Giardia***

An adapted Luria-Delbrück fluctuation tests^11^ was designed to determine the *Giardia* switching rates in the presence or absence of the anti-VSP antibodies. Briefly, 10 trophozoites (N0) of *Giardia* clones expressing VSP417 were used to initiate 12 independent cultures of 1 ml each (C=12) and cultured for additional 72 h in the presence of anti-VSP417 mAb 7C2 (n=6) or of the unrelated mAb 8F12 (n=6) at 50 nM. Subsequently, cells from each well were counted (Nt), washed and resuspended in 2.4 ml of culture medium without mAbs and plated into 12 new wells. After 24 h, trophozoites were collected from all wells and subjected to IFAs using Alexa Fluor 488-conjugated anti-VSP mAbs 7C2 and 1,000 parasites per well were counted to record positive (non-switchers) and negative (switchers) cells. Then, the P0 method^11,12^ was used to calculate the VSP switching rates (µ), where P0 is the fraction of non-switchers per culture. The number of switchers per culture (m) was considered as the -ln P0. The switching rate was calculated as m divided by the number of cells per culture at 72 h of incubation with the antibodies (Nt). The mean and the variance in the number of switchers were also determined. When the variance and the mean are similar, their ratio represents inducible cultures, whereas higher values corresponds to spontaneous switching^11,12^.

***In vitro* switching dynamics**

*G. lamblia* trophozoites of clone VSP417 were grown in TYI-S-33 medium with 50 nM of mAb 7C2 or the unrelated mAb in 8-ml culture tubes. The initial number of trophozoites used for each chosen time point was calculated to reach a final number of 3x10^6^ cells after the stipulated culture time, assuming a generation time of 8 h, to ensure exponential growth. At each time point, trophozoites were counted and treated with Alexa Fluor^TM^-488 direct-labelled 7C2 mAb (Alexa Fluor^TM^-488 labelling kit; Invitrogen^TM^, Cat. # A10235) and VSP417^(-)^-pAb in TYI-S-33 medium on ice for 40 min. The cells were then washed twice with ice-cold phosphate-buffered saline (PBS) and fixed with 4% formaldehyde (PFA) in sodium phosphate 0.1 M pH 7.2 for 1 h. Then, cells were washed twice with PBS and treated with ammonium chloride 50 mM for 10 min and with 1% BSA in PBS (blocking solution) for 15 min. Labelling for cell cytometry was performed using goat anti-mouse IgG (H-L)-PE (Invitrogen^TM^, Cat. # PA1-84395, 1/6000), goat anti-rat IgG (H-L)-biotin 1/2000 (Invitrogen^TM^, Cat. # A18869), and streptavidin-Alexa Fluor^TM^ 488 (Invitrogen^TM^, Cat. # S11223, 1/6000) in blocking solution. Super-resolution structured illumination microscopy (SR-SIM) analysis was performed on trophozoites at 24 h post induction. Fixed trophozoites were attached to a coverslip using poly-L-lysine solution 0.1% (Sigma-Aldrich, Cat. # P8920), blocked and treated with goat anti mouse IgG (H+L)-Alexa Fluor^TM^ 488 (Invitrogen^TM^, Cat. # A11001) and goat anti-rat IgG (H-L)-Alexa Fluor^TM^ 546 (Invitrogen^TM^, Cat. # A11081) secondary antibodies at a dilution of 1/200 in blocking solution for 1 h. DNA staining was performed with DAPI. Images were taken using a Zeiss Elyra PS1 microscope system. Images were acquired with five grid rotations and analysed with the ZEN software (Zeiss).

***In vivo* switching dynamics**

Gerbils were infected by orogastric inoculation of 2x10^5^ trophozoites of clone VSP417 resuspended in 0.5 ml of PBS. Randomly selected gerbils were euthanized on days 6, 9, 10, 11, 12, 13 and 14 post-infection and the first 14-cm portion of the small intestine was isolated and dissected longitudinally before being incubated on ice-cold PBS for 30 min. The supernatants were collected and *Giardia* trophozoites were quantified in a haemocytometer and VSP characterization was carried out by flow cytometry, as explained above.

**Cloning and transfection**

The plasmid pTUBpac^13^ was used to express cytoplasmic tail-variants of VSPH7. To construct the different VSPH7 variants, the same forward primer (fwd, 5’-CAT GCC ATG GAT GTT TCT ATT AAT TAA TTG CCT A-3’) was used in combination with different reverse primers. VSPH7WT (wt_rev, 5’-GCG GAT ATC CGC CTT CCC GCG GCA GAC GAA-3’) is the wild-type version of the VSPH7 (CRGKA); VSPH7AxK (ak_rev, 5’-GCG GAT ATC CGC CGC CCC GCG GCA GAC GAA-3’) has Ala instead of a Lys in the tail. Similar modifications were made in the variants VSPH7AxR (ar_rev, 5’-GCG GAT ATC CGC CTT CCC CGC GCA GAC GAA-3’) and VSPH7KxC (kc_rev, 5’-GCG GAT ATC CGC CTT CCC GCG CTT GAC GAA-3’), where Arg was replaced with Ala and Cys was replaced with Lys, respectively. For TMD variants, the pTUBH7HApac was modified for the repositioning of the ApaI sequence from the 5´end of the VSPH7 to a new position upstream of the TMD sequence. PCR mutagenesis by overlap extension was performed using the following primers: Fwd_A, 5’-GAA CCA TGG GGC TCT TAA TTA ATT G-3’; Rev_B, 5’-GAG GAG AGG TTG GGC CCA CTA-3’; Fwd_C, 5’-TAG TGG GCC CAA CCT CTC CTC-3’; and Rev_D, 5’-AAT TCA CTG CGG CCG CAA CTC-3’. The vector obtained was named pTUBH7ApaI. Then, two VSPH7 TMDs variants were obtained through the annealing of the following primers: Fwd_GAS/M, 5’- C AAC CTC TCC TCT ATG GCG ATC GCA ATG ATC TCG GTG ATG GTC ATT GTC GTC GTC ATG GGC CTC GTC ATG TTC CTC TGC TGG TGG TTC GTC TGC CGC GGG AAG GCG TGA GC -3’; Rev_GAS/M, 5’- GGC CGC TCA CGC CTT CCC GCG GCA GAC GAA CCA CCA GCA GAG GAA AAG GAC GAG GCC AAG GAC GAC GAC AAT GAC GGC CAC CGA GAT AAG TGC GAT CGC AAG AGA GGA GAG GTT GGG CC-3’; Fwd_Aro/V, 5’-CAA CCT CTC CTC TGG CGC GAT CGC AGG CAT CTC GGT GGC CGT CAT TGT CGT CGT CGG AGG CCT CGT CGG CGT TCT CTG CGT TGT TGT TGT CTG CCG CGG GAA GGC GTG AGC-3’; and Rev_Aro/V, 5’-GGC CGC TCA CGC CTT CCC GCG GCA GAC AAC AAC AAC GCA GAG AAC GCC GAC GAG GCC TCC GAC GAC GAC AAT GAC GGC CAC CGA GAT GCC TGC GAT CGC GCC AGA GGA GAG GTT GGG CC-3’. In the GAS/L TMD, Gly residues were replaced with Met, and in the ARO TMD, Trp and Phe residues were replaced with Val. For the introduction of TMD variants into the pTUBH7ApaI vector, both the vector and TMDs were digested using ApaI and NotI and ligated. Deletion and mutations were confirmed by sequencing using dye terminator cycle sequencing (Beckman Coulter). Trophozoites were transfected by electroporation and selected with puromycin (InvivoGen, Cat. # Ant-pr-5), as previously described^14^.

***In silico* VSP TMD oligomerization models**

Oligomerization capability of the TMD was analysed with THOIPA^15^ and PREDIMMER^16^ and visualized with VMD^17^. Alignments were performed with Clustal Omega^18^ and WebLogo^19^. Figures and graphic representations were created with BioRender.com.

**Isolation of detergent-resistant membranes (DRMs)**

DRMs were prepared as previously described^20^, with modifications. Trophozoites (2.5x10^7^) of clone VSP417 were resuspended in 5 ml of TYI-S-33 medium with 50 nM of mAb 7C2 or without mAb at 37°C for 30 min. Pre-treatment with 10 mM methyl-β-cyclodextrin (MβCD; Sigma-Aldrich Cat. # 128446-36-6) was performed in PBS supplemented with ascorbic acid (0.01% w/v) and cysteine (2% w/v), pH 7.2 (PBSAC) for 30 min and then incubated with 50 nM of mAb 7C2 in PBSAC. Trophozoites were then washed with TNE buffer (50 mM Tris-HCl pH 7.5; 150 mM NaCl; 1 mM EDTA) and lysed in 500 µl of TNE buffer supplemented with 1% w/v Triton X-100 and 2X cOmplete™ protease inhibitor cocktail (Merck^®^, Cat. # 11836145001) on ice for 30 min. An equal volume of ice-cold TNE 85% sucrose was added to cell lysates to adjust sucrose concentration at 42.5%. The lysates were then placed at the bottom of a 13.2-ml ultra-clear centrifuge tube (Beckman) and carefully overlaid with 5 ml of 35% sucrose in TNE and 1 ml of 5% sucrose in TNE. Tubes were centrifuged at 250,000 g in a SW41Ti rotor (Beckman Coulter) at 2°C for 20 h. Fourteen fractions of 500 µl were carefully obtained from the top of each tube and 36 µl of each fraction was subjected to SDS-PAGE under non-reducing conditions^19^. VSP417 was detected by Western blotting using mAb 7C2 (1:1,000) and peroxidase-conjugated goat anti-mouse IgG (H-L) (Invitrogen^TM^, Cat. # 626520; 1:5000). Fractions 3-12 and 13-14 were considered DRM and DSM, respectively.

**Dynamics of antibody-bound VSPs elimination**

For electron microscopy, 5x10^6^ trophozoites of clones VSP417, VSP1267 and VSPH7 were treated with 2.5 nM of their cognate anti-VSP mAbs in TYI-S-33 medium on ice for 1 h. Then, cells were washed three times with ice-cold PBS, resuspended in 7 ml of TYI-S-33 medium and incubated at 37°C for 0, 15, 30 and 60 min. At each time point, trophozoites were placed on ice for detachment, washed twice with ice-cold filtered PBS, and fixed with 4% FA in sodium phosphate 0.1 M pH 7.2 for 1 h. Cells were washed twice with PBS and treated with ammonium chloride 50 mM for 10 min and with blocking solution for 15 min. Immunogold labelling was performed by treating the trophozoites with goat anti-mouse IgG-(H-L)-gold 10 nm (Abcam, Cat. # ab39619) in a dilution of 1/10 in blocking solution for 2 h and fixed again with 2.5% glutaraldehyde in 0.1 M cacodylate buffer (pH 7.2).

**Scanning electron microscopy (SEM) and Helium ion microscopy (HM)**

Briefly, glutaraldehyde-fixed cells were post-fixed in 1% OsO_4_ for 15 min. Then, samples were dehydrated in crescent series of ethanol up to 100%, critical point-dried with liquid CO_2_ and sputter-coated with carbon in order to observe the cell surface in detail. The specimens were examined in a Quanta^TM^ SEM (FEI Co., The Netherlands) equipped with FEG filament. Images were obtained via secondary electron (SE) and/or backscattered electron (BSE) detection at an accelerating voltage of 15 kV. For High-Resolution Scanning Microscopy analysis, the sputter-coating step was performed using a thin layer (2 nm) of platinum, and cells were observed on an Auriga^TM^ High-Resolution SEM (Zeiss). For HM, cells were processed as described above and observed, without any subsequent coating, under a Zeiss Orion Helium Ion Microscope.

**Transmission electron microscopy (TEM)**

Glutaraldehyde-fixed cells were post-fixed with 1% OsO_4_ and 0.8% potassium ferrocyanide for 40 min. The samples were dehydrated in crescent grades of acetone up to 100% and embedded in epoxy resin. Ultrathin sections (50-60 nm thick) were cut, collected and stained with uranyl acetate and lead citrate. Lastly, the samples were analysed using a Tecnai^TM^ Spirit TEM (FEI Co.).

**Microvesicle purification and validation**

Exponentially growing trophozoites (1.5x10^8^) of clone VSP417 were washed twice with filtered PBS at 37°C and incubated 4 h at 37 °C in the absence or the presence of mAb 7C2 (50 nM) in microvesicle purification medium (ultrafiltrated TYI-S-33 medium, 100 kDa MWCO, supplemented with 3% adult bovine serum previously ultracentrifuged overnight at 250,000 g). Tubes were ice cooled and then centrifuged in a SW41Ti rotor (Beckman) at 4°C for 10 min. Supernatants were collected in 50-ml centrifuge tubes and centrifuged at 3,000 g at 4°C for 40 min; this step was repeated once. Then, the supernatants were concentrated 10X in a centrifugal filter device (Centricon^TM^ Plus-70-100K, Millipore^®^, Cat. # UFC710008) and ultracentrifuged in a Beckman^®^ SW41Ti rotor (25,000 rpm, 2 h, at 4°C). Pellets were washed in an equal amount of filtered ice-cold PBS and ultracentrifuged again to recover the MVs. In the absence of mAb, no MVs were detected and, consequently, untreated samples were discarded for subsequent analysis. Purification of MVs was validated by TEM. Briefly, a pellet of microvesicles was resuspended in 500 µl of 1% BSA in PBS with anti-mouse IgG-Gold 10 nm (Abcam, Cat. # ab39619; 1:100) and incubated on ice for 2 h. Next, the samples were washed twice with filtered PBS and fixed with 2% PFA in PBS. Microvesicles without gold labelling were also fixed. To obtain TEM images, a Formvar-coated nickel grids were floated on 10 µl of microvesicles suspension for 20 min. Subsequently, microvesicles adsorbed on the grids were post fixed in 1% glutaraldehyde. The grids were rinsed with dH_2_O and contrasted successively in 2% uranyl acetate pH 7 and 2% methylcellulose/0.4% uranyl acetate, pH 4. Microvesicles were visualized using a Leo 906-E (Zeiss) transmission electron microscope.

**Proteomics of purified microvesicles**

Monoclonal antibody 7C2-induced microvesicles from three independent *Giardia* clones expressing VSP417 and treated with the mAb 7C2 were analysed with MS/MS. Purified microvesicles were lysed in RIPA buffer with cOmplete^™^ protease inhibitor cocktail (Merck^®^, Cat. # 11836145001). Fifteen µg of each sample were run 1 cm in SDS-PAGE under non-reducing conditions. The following procedure and analysis were done by MSBioworks.com: In-gel digestion was then performed on each sample using a robot (ProGest, DigiLab) with the following protocol: washed with 25 mM ammonium bicarbonate followed by acetonitrile, reduced with 10 mM dithiothreitol at 60°C followed by alkylation with 50 mM iodoacetamide, digested with sequencing grade trypsin (Promega) at 37°C for 4 h and quenched with formic acid. Half of each digested sample was analysed by nano LC-MS/MS with a Waters NanoAcquity HPLC system interfaced to a Thermo Fisher Q Exactive mass spectrometer. Peptides were loaded on a trapping column and eluted over a 75-μm analytical column at 350 nL/min; both columns were packed with Luna C18 resin (Phenomenex). Total instrument time used was 2 h. The mass spectrometer was operated in data-dependent mode, with the Orbitrap operating at 70,000 FWHM and 17,500 FWHM for MS and MS/MS, respectively. The 15 most abundant ions were selected for MS/MS. Data were processed using both Mascot (Matrix Science) with the following parameters: Enzyme, Trypsin/P; Database, NCBI *Giardia lamblia* RefSeq 2.1 proteome (concatenated forward and reverse plus common contaminants); Fixed modification, Carbamidomethyl (C); Variable modifications, Oxidation (M), Acetyl (N-term), Pyro-Glu (N-term Q), Deamidation (N/Q); Mass values, Monoisotopic; Peptide Mass Tolerance, 10 ppm; Fragment Mass Tolerance, 0.02 Da; Max Missed Cleavages, 2. Mascot DAT files were parsed into Scaffold (Proteome Software) for validation, filtering and to create a non-redundant list per sample. Data were filtered using 1% protein and peptide FDR and requiring at least two unique peptides per protein. The mass spectrometry proteomics data have been deposited to the ProteomeXchange Consortium via the PRIDE partner repository^21^ with the dataset identifier PXD031141 and 10.6019/PXD031141 (Reviewer’s account details: Username**:**; Password**:** QmM5xe8V).

**Calcium treatments and quantification**

The involvement of calcium was determined by incubating 1-2x10^4^ trophozoites of clone VSP417 in PBS and stimulated in culture medium with different concentrations of CaCl_2_ (0.1-10 mM) at 37°C for 72 h in the presence or absence of their cognate anti-VSP mAb; then, the percentage of the former VSP was determined in the parasite population by IFA. On the other hand, the effect of the intracellular calcium chelator during AV induced by antibodies was studied. *Giardia* trophozoites (1 x 10^6^) of clone VSP417 were incubated in culture medium supplemented with 100, 250 or 500 µM of BAPTA-AM (Sigma-Aldrich, Cat. # A1076) at 37°C for 72 h, in the presence of 50 nM of mAb 7C2 or an unrelated antibody. After that period, parasites were washed twice in cold PBS, collected by centrifugation, and analysed using IFA. For intracellular calcium measurements, Fluo 4-AM was freshly prepared in dehydrated DMSO before each experiment. Trophozoites expressing VSP417 were harvested by centrifugation (1,000 g for 5 min) and the cells were then resuspended in Hanks’s balanced salt solution (HBSS) supplemented with 10 mM HEPES. Cells were then incubated with Fluo 4-AM (10 µM) and Pluronic F-127 (0.04%) at room temperature for 30 min and then washed three times for at least 15 min to allow complete intracellular de-esterification of the dye. Finally, trophozoites were incubated 30 min at 37°C in HBSS-HEPES. Kinetic experiments were performed with a Cary Eclipse spectrofluorimeter (Agilent Technologies^®^) equipped with a stirred cuvette holder. A 0.3 mm path cuvette was used. Trophozoites were excited at 495 nm and emission at 520 nm with excitation/emission slit widths of 5/5 nm.

**Extraction of total RNA from bloodstream forms of *T. brucei***

Cultures were amplified in HMI-9 medium supplemented with 20% heat-inactivated foetal bovine serum (Gibco, Cat. # A3840301). Subsequently, the parasites (between 1x10^7^ and 2x10^7^) were lysed in 0.5 ml of Trizol (Invitrogen^TM^, Cat # 15596026) and total RNA was extracted according to the manufacturer's protocol. The RNA obtained was dissolved in 45 µl of DEPC water and treated with 2 U of DNAse I in a final volume of 50 µl at 37°C for 1 h. Subsequently, a second RNA purification was carried out by adding 150 µl of DEPC water and 200 µl of chloroform. The mixture was vortexed and centrifuged at maximum speed for 30 seconds. Then, the upper aqueous phase was collected and 70 µl of 3 M potassium acetate pH 5.2 and then 100 µl of absolute ethanol were added. The mixture was incubated at -20°C for 15 min and centrifuged at 12,000g at 4°C for 30 min. The pellet obtained was washed twice with 200 ml of cold 75% ethanol and subsequently left to dry for 10 min. Finally, the pellet was dissolved in 36 ml of DEPC water and incubated in a thermostatic bath at 55°C for 10 min. The integrity of the obtained RNA was confirmed by agarose gel electrophoresis. The purified RNA was stored at -70°C.

**RT-PCR for amplification of VSG transcripts**

The reverse transcription was performed using 1 µg of total RNA. ProtoScriptII^TM^ reverse transcriptase (200 U) (NEB^®^, Cat # M0368) was used according to the manufacturer's instructions using random hexamer primers. The cDNA was stored at -20°C. Controls were carried out in parallel without the addition of reverse transcriptase to rule out possible contaminations with genomic DNA. For the amplification of the transcripts encoding VSGs, a PCR was performed with the generic primers Splice_leader and VSGall, whose sequences are Splice_leader 5’-AGT TTC TGT ACT AT-3’, VSGall 5’-GTG TTA AAA TAT ATC-3’. For the PCR mix, 2.5 U of Taq polymerase, 1X Taq Buffer, 1.5 mM MgCl_2_, 0.2 mM dNTPs, 0.6 µM of each primer and 2 µl of cDNA (or control without reverse transcription) were used in a final volume of 50 µl. The PCR consisted of 5 min incubation at 94°C, followed by 30 cycles of denaturation (30 seconds at 94°C), hybridization (30 seconds at 40°C) and extension (2 min at 72°C) and an incubation of 10 min at 72°C. PCR products with a size between 1.5 and 2 kbp were purified from agarose gels and submitted to Macrogen (South Korea) for sequencing using the Splice_leader and VSGall primers.

**Switching rate determination in *T. brucei***

A similar Luria-Delbrück fluctuation test used for *Giardia* was designed to determine the switching rates in *Trypanosoma* *brucei* but using two different methods for the selection of switchers: anti-VSG pAb at cytotoxic concentrations or *VSG* RNAi. In the first case, 18 cultures of polymorphic bloodstream forms of *T. brucei* expressing VSG AnTat1.1 were initiated using ~20 parasites per well (N0) and grown for 72 h in the presence of a sub-lethal concentration of the anti-VSG pAb (1/1,000) (n=9) or of preimmune serum (1/50) (n=9). After this period, cells from each well were counted (Nt) and plated into 32 new wells supplemented with a lethal concentration of the anti-VSG AnTat1.1 antibody (1/50) for 4 days to kill non-switchers. The wells containing live parasites were subjected to cDNA sequencing to determine the identity of the expressed VSGs and to determine the possible presence of non-switchers. In the case of selection of switchers by *VSG* RNAi, 16 independent 1 ml cultures of *T. brucei* EATRO1125 *VSG* AnTat1.1 RNAi were initiated at concentrations ranging from 5 to 50 cells ml^−1^ (N0). Cells were amplified for typically 8 or 9 generations (Nt) in the presence of a sub-lethal concentration of the anti-VSG pAb (1/1,000) or of preimmune serum. Then, *VSG* RNAi was induced by adding doxycycline (1 mg ml^−1^). Each culture was spread over 10 wells of a 96-well plate and wells were allowed to grow for 6–8 days before scoring. The total number of wells were recorded as either switchers or non-switchers. As in *Giardia*, the P0 method^11,12^ was used to determine *T. brucei* switching rates.

**Statistical analyses**

No statistical methods were used to predetermine sample size, except for the Luria-Delbrück fluctuation tests. The experiments were randomized. The investigators were blinded to allocation during experiments and outcome assessment. Average (mean) and s.e.m. were calculated in Excel. Statistical significance is based one-way or two-way ANOVA on datasets with Dunnett's and Sidak's multiple comparisons test or Bonferroni post-test, respectively, using GraphPad Software (Prism). All figures show the mean value of three independent experiments ± s.e.m. Statistically significant differences are indicated in each graph as ^*^*p*< 0.05, ^**^*p*< 0.01, ^***^*p*< 0.01, ^****^*p*< 0.001 and ns=not significant.

**Methods References**

**Extended Data Tables and Figures**

**Extended Data Table 1| Murine monoclonal antibodies against different *Giardia* VSPs and CWP1.** Monoclonal antibodies (mAbs) specific for a given VSP of assemblages A1 (WB isolate) or B (GS/M-83 isolate). For VSP1267, two different mAbs were tested (IgG_1_ and IgM). The mAb against Cyst Wall Protein 1 (CWP1; a *Giardia* protein not expressed in proliferating trophozoites) was used as control.

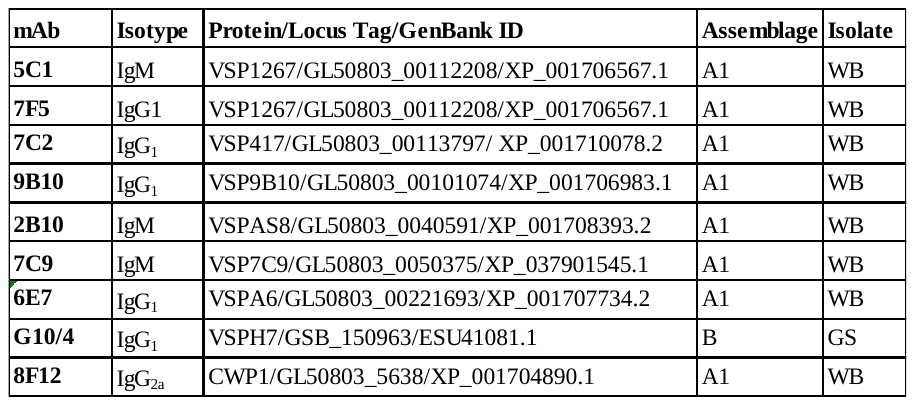

**Extended Data Table 2| Expression of novel VSPs after mAb-induced antigenic switching.** Two 96-well plates were used to distribute 0.5 cells per well of clone VSP417. After 5 days of culture in the presence of mAb 7C2 (50 nM), the composition of *Giardia* populations for 20 randomly selected wells was determined by IFA using mAbs against different VSPs. Values are percentages of positivity to the corresponding anti-VSP mAb.

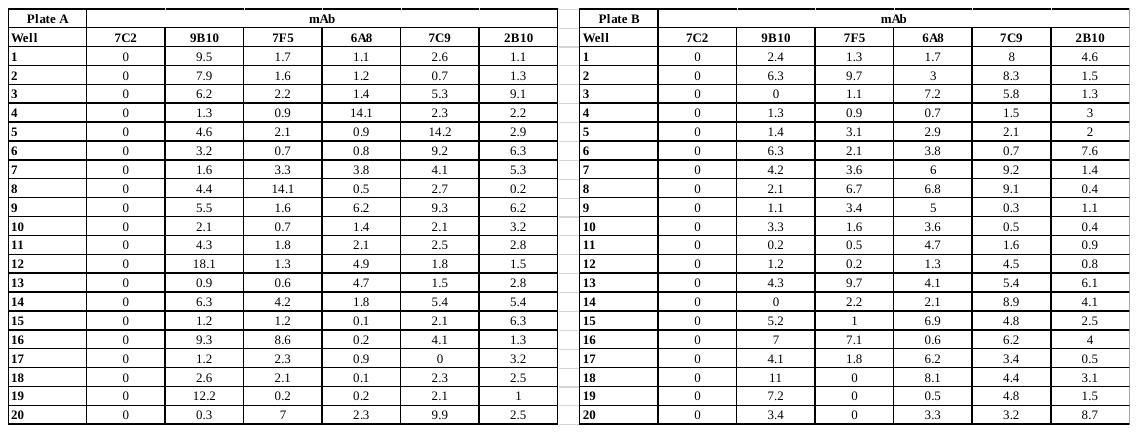

**Extended Data Table 3| Adapted Luria-Delbrück Fluctuation Test in *Giardia*.** Trophozoites of clone VSP417 were culture for 72 in the absence (control unrelated mAb; Exp. 1-6) or the presence of the anti-VSP417 mAb 7C2 (50 nM) (+ mAb; Exp. 7-12) and subjected to the adapted Luria-Delbrück fluctuation test. Incubation with the mAb directed to the cognate VSP increased the switching rate (µ) by ~72 times as compared to the controls treated with the unrelated mAb. C represents the total number of independent cultures, N0 is the number of parasites in the initial inoculum, Nt indicates the number of parasites at 72 h, P0 is the fraction of non-switchers, m is the –ln P0. Similar variance (Var.) and mean values indicates that the increase in µ is induced by the treatment.

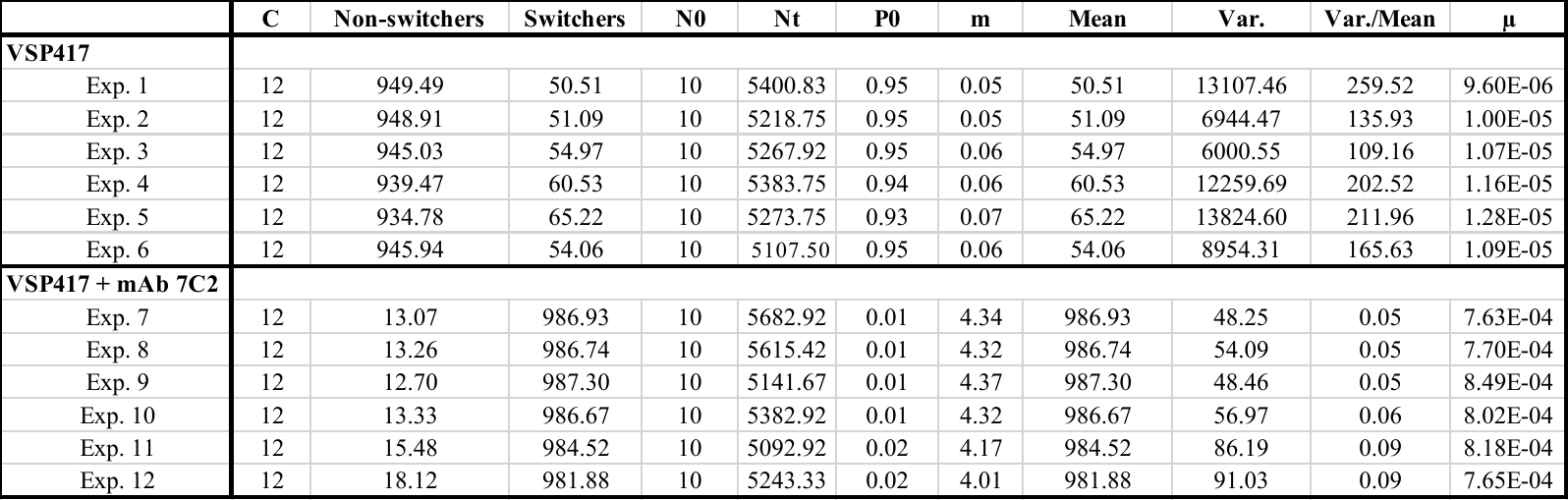

**Extended Data Table 4| Proteomic analysis of purified microvesicles**. Purified MVs obtained after 4 h of incubation with mAb 7C2 of trophozoites expressing VSP417 were subjected to proteomics analysis as described in Methods. Columns indicate the GenBank™ accession number, the locus tag, the original name of the proteins and a newly proposed one, the total number of identified peptides (SpC), the SpC of each independent experiments, the presence of absence of a TMD and/or a signal peptide (SP) and comments about the predicted subcellular localizations of all detected proteins are shown. VSPs are highlighted in yellow, α-giardins (annexins) in light blue and ALIX/PDCDIP in green. (See separate Excel table)

**Extended Data Table 5| Adapted Luria-Delbrück Fluctuation Tests in pleomorphic bloodstream forms of *T. brucei* using different selection methods. a,** Selection of switchers by *VSG* RNAi. Independent cultures were initiated with 5 and 50 cell/ml (N0) of *T. brucei* clone EATRO1125 *VSG* AnTat1.1 RNAi expressing VSG AnTat1.1 were grown for 8 to 9 generations (Nt) in the absence (control preimnune sera) or presence (+ pAb) of sub-lethal dilutions (1/1,000) of the specific anti-VSG antibody. After addition of doxycycline, cultures were plated and grown for 6-8 days. Wells containing the surviving switchers were identified and values subjected to the adapted Luria-Delbrück fluctuation test. **b**, Selection of switchers by high antibody concentrations**.** Approximately 20 cells (N0) per culture (n=18) of *Trypanosoma brucei* clone EATRO1125 expressing VSG AnTat1.1 were grown for 8 to 9 generations either in the absence (control preimnune sera; Exp. 1) or presence of sub-lethal dilutions (1/1,000) of the specific anti-VSG antibody (+ pAb; Exp. 2). After addition of a lethal dilution of the anti-VSG pAb (1/50), cultures were plated into 32 new wells and growth for 72 h. The surviving switchers were identified and values subjected to the adapted Luria-Delbrück fluctuation test. In both cases, the estimated switching rate (µ) was increased in cells confronted to the antibody against VSG AnTat1.1. N0 is the number of parasites in the initial inoculum, Nt indicates the final number of parasites before selection, P0 is the rate of non-switchers, m is the –ln P0. Similar variance (Var.) and mean values indicates that the increase in µ is induced by the treatment with the anti-VSG pAb.

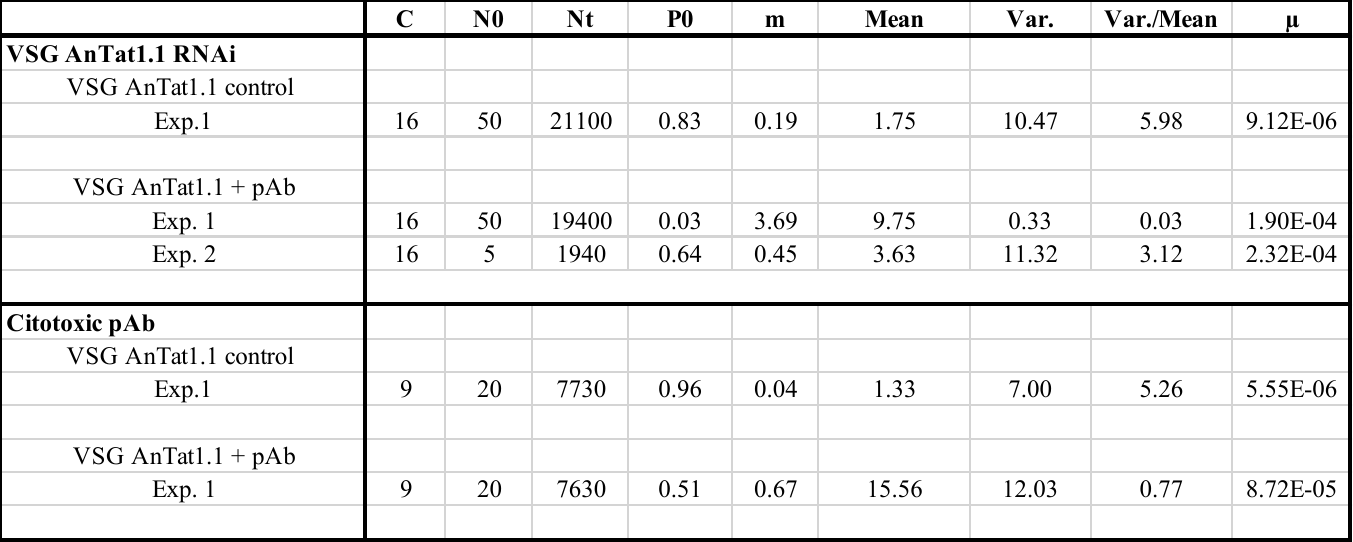

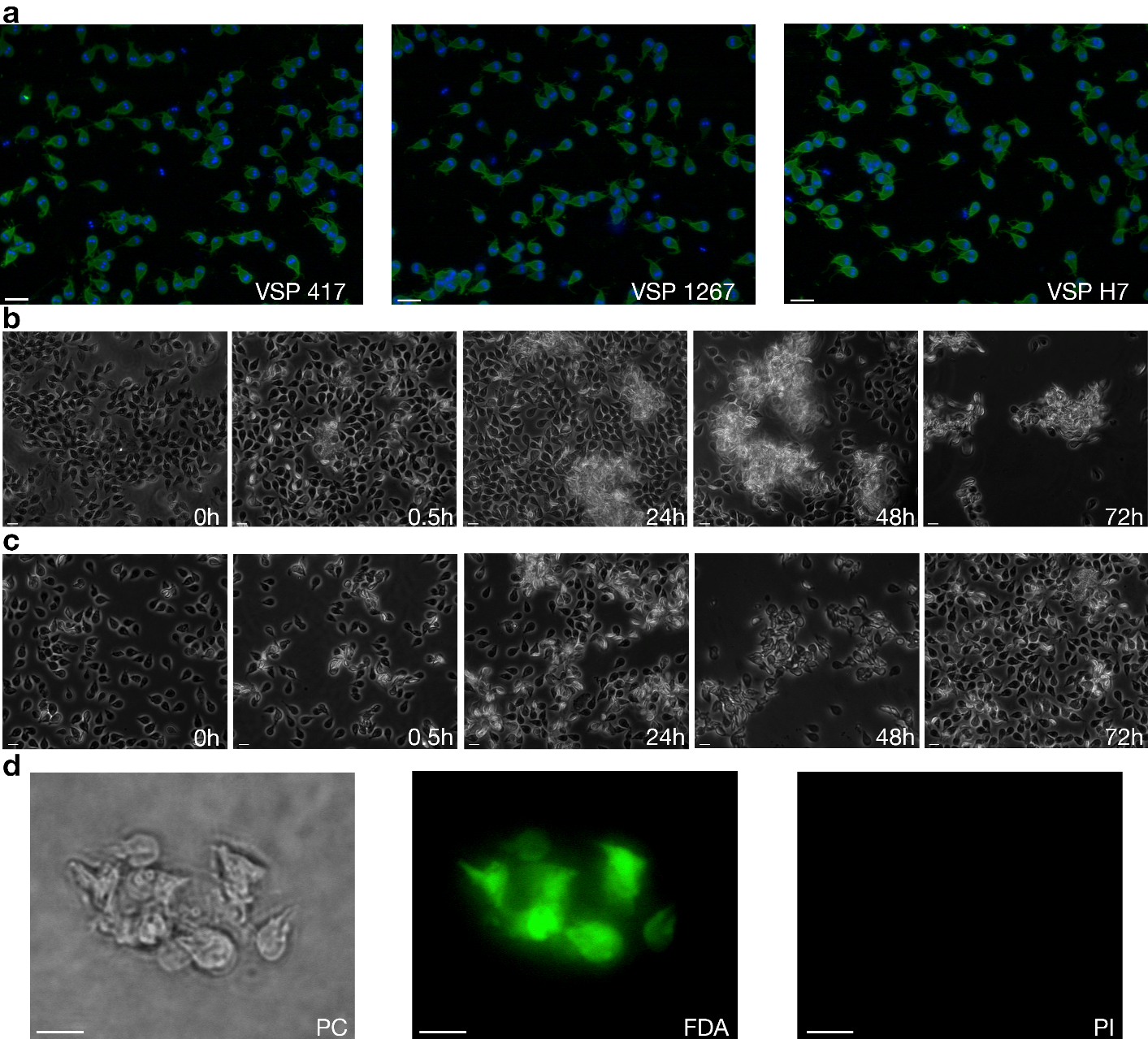

**Extended Data Figure 1| Effect of antibodies on *Giardia* trophozoites. a,** Representative immunofluorescence images of the different VSP clones used (VSP, green; Nuclei, blue). **b**, *Giardia* trophozoites of clone VSP417 incubated in the presence of 100 µM of mAb 7C2. A rapid agglutination of the cells was observed and clusters of cells remained grouped but alive at 72 h of culture with mAb 7C2. **c**, *Giardia* trophozoites of clone VSP417 incubated in the presence of 50 nM of mAb 7C2. A rapid agglutination of the cells was observed, but cells then reattached to the glass tube at 72 h of culture with mAb 7C2. **d**, Representative image of a cluster of *Giardia* trophozoites expressing VSP417, incubated for 48 h in the presence of mAb 7C2, and stained with FDA to label live cells (green) and with PI to label the nuclei of dead cells (red). All trophozoites of the aggregate are alive after that period in the presence of the anti-VSP antibody. Scale bars 10 µm.

| **Expression of the original VSP** | **Original VSP / number of clones** |
| --- | --- |
| Clone VSPH7 control | 31/31 |
| Clone VSP1267 control | 27/28 |
| Clone VSP417 control | 27/28 |
| Clone VSPH7 Exp. 1 | 1/27 |
| Clone VSPH7 Exp. 2 | 0/29 |
| Clone VSPH7 Exp. 3 | 1/31 |
| Clone VSP1267 Exp. 1 | 1/33 |
| Clone VSP1267 Exp. 2 | 0/35 |
| Clone VSP1267 Exp. 3 | 0/30 |
| Clone VSP417 Exp. 1 | 0/26 |
| Clone VSP417 Exp. 2 | 1/37 |
| Clone VSP417 Exp. 3 | 0/35 |

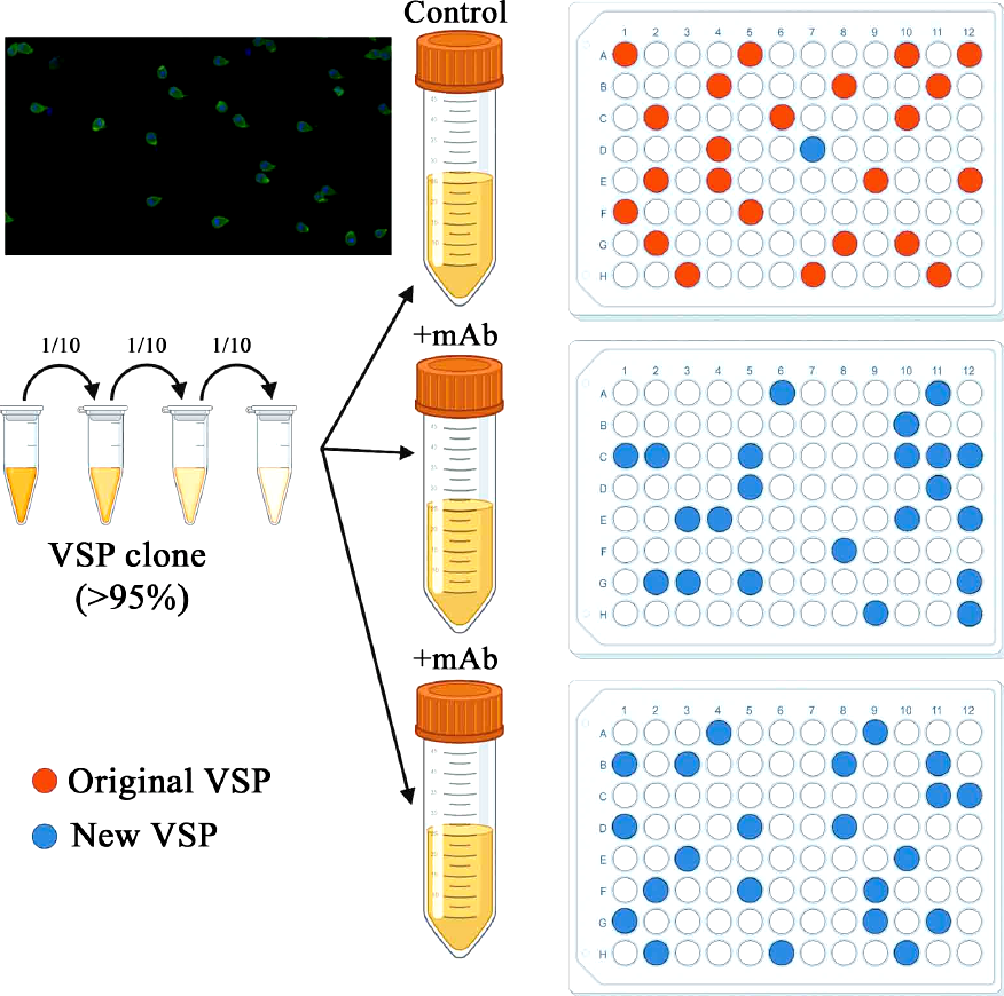

**Extended Data Figure 2| Cytotoxicity and switching assay. Left.** Schematic representation of the in-cell antigenic switching assay (**left**). *Giardia* trophozoites expressing a particular VSP (almost 99 positive by IFA; upper left corner) were subjected to limiting dilution to reach 50 trophozoites/ml and subsequently distributed in 96-well plates containing culture medium including either an unrelated mAb (mAb 8F12; top plate) or anti-VSP mAb (+mAb) at a final concentration of 50 nM. After 5 days of culture, the total number of growing clones and their reactivity to the mAb was determined. Orange wells represent clones expressing the original VSP and blue wells represent clones expressing a different VSP. **Right**, Table showing the number of clones expressing the original VSP vs the total number of growing clones in three independent experiments. In the presence of the corresponding mAb, most clones expressed a different VSP after the treatment, demonstrating that anti-VSP antibodies induce antigenic variation.

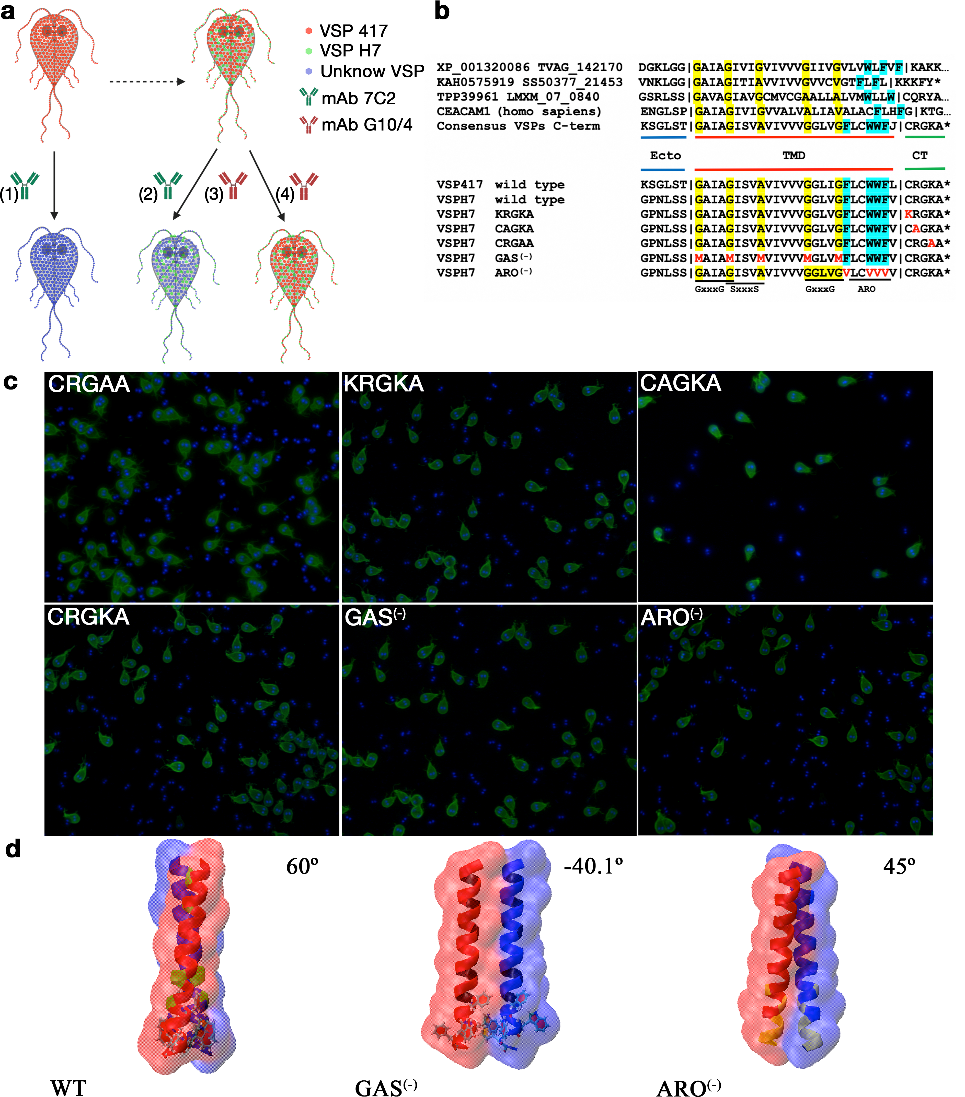

**Extended Data Figure 3| The TMD of VSPs is involved in VSP clustering induced by anti-VSP antibodies. a**, Schematic representation of the experimental design for evaluating the involvement of the CT and the TMD of VSPs during antigenic switching. The diagram shows the reporter VSP417 in red (endogenously expressed) and the VSPH7 in green (constitutively expressed), the latter carrying different mutations in the C-terminal conserved region that includes the TMD and the CT (see B). Possibility 1 shows the induction of AV by antibodies to the endogenous VSP417. Possibilities 2-4 depict the different outcomes expected when parasites co-expressing two VSPs (VSP417 in red and VSPH7 in green) are exposed to either an anti-VSP417 or an anti-VSPH7 mAb. The induction of AV of the reporter VSP417 (option 3) or not (option 4), reflecting the involvement or not of the conserved C-terminus of VSPs as transducer of the antibody signal. **b**, Alignment of the C-terminal sequences of *Trichomonas vaginalis* G3 Hypothetical Protein XP_001320086 (V), *Spironucleus salmonicida* Cysteine-rich membrane protein 2 KAH0575919, *Leishmania Mexicana* Hypothetical protein CGC21_25575, human CEACAM1 and the consensus sequence of VSPs (top). Different variants of VSPH7 were designed to disrupt the GAS motifs (GAS^(-)^) and the motif rich in aromatic amino acids (ARO^(-)^). **c**, *In vivo* labelling of trophozoites using mAb G10/4 (left panel) showed surface localization of each VSPH7 variant. Scale bar, 10 µm. **d**, Modelling of the wild type (WT), GAS^(-)^ and ARO^(-)^ variants of the TMD of VSPH7 showing the differences in oligomerization potential of the mutants compared to the WT TMD (Rosetta ddG: WT -51.94 > ARO^(-)^ -35.59 > GAS^(-)^ -21.86).

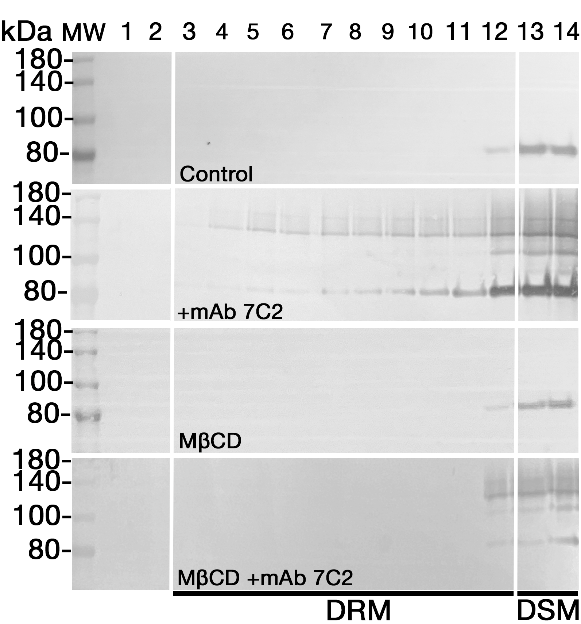

**Extended Data Figure 4| The VSP/anti-VSP complex segregates into lipid raft-like microdomains.** Western blotting analysis of the distribution of VSP417 into detergent resistant membranes (DRM) preparations compared to its presence in detergent soluble membranes (DSM) after treatment with 50 nM of mAb 7C2 (+mAb) or the unrelated mAb 8F12 for 1 h. Numbers on the top represent the fourteen collected fractions. Pre-treatment of the cells with methyl-β-cyclodextrin (10 mM) before incubation with mAb 7C2 or a control antibody are also shown (MβCD+mAb). Although at 1 h most of the VSP/antibody complex remains in the detergent soluble membrane (DRM) fractions, the complex redistributes into the detergent resistant membrane (DRM) fractions upon treatment with the antibody. This redistribution is abolished if the trophozoites were previously treated with 10 mM of the cholesterol-rich disrupting domains MβCD.

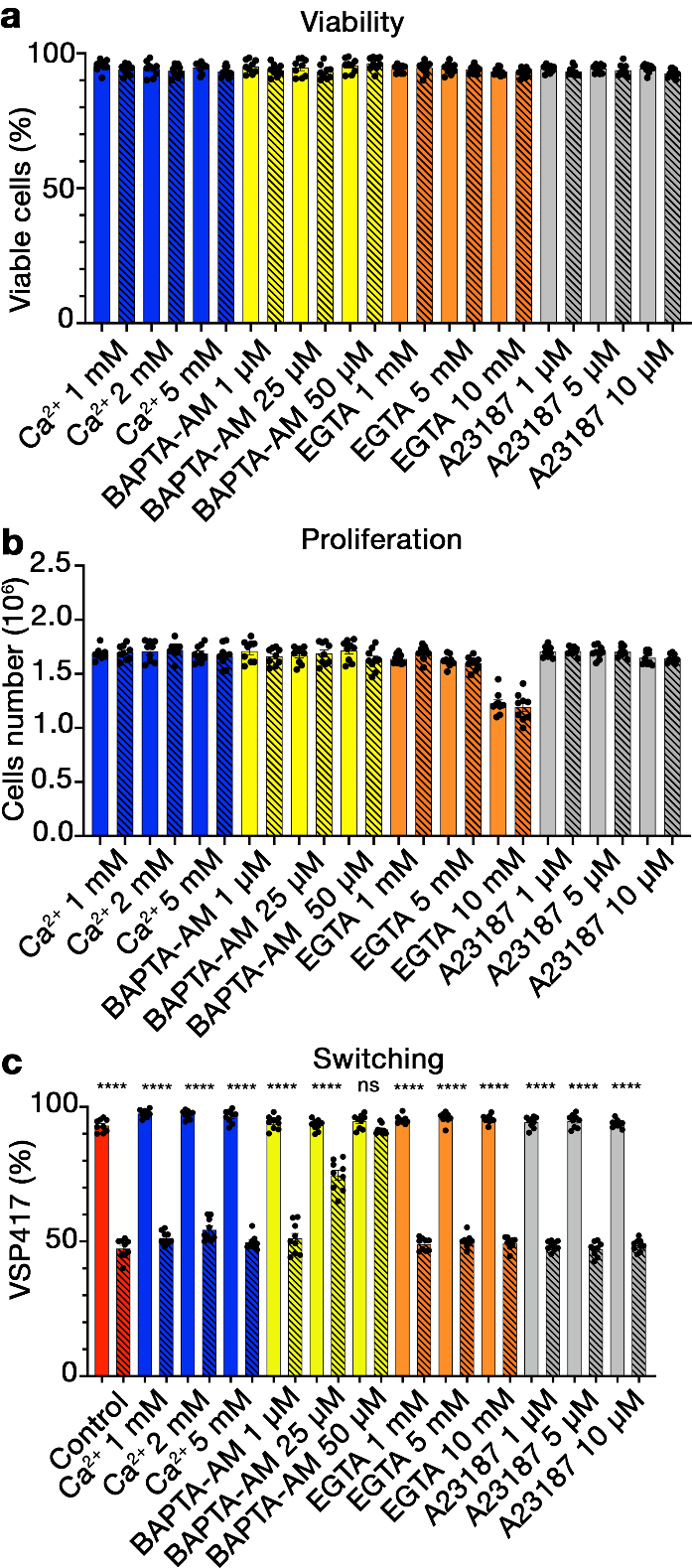

**Extended Data Figure 5| Effect of different Ca^2+^ treatments on viability, proliferation and VSP switching.** VSP417-expressing trophozoites were grown in the presence of a control mAb (solid bars) or of mAb 7C2 (striped bars) for 72 h. The effect of different concentrations of extracellular calcium, EGTA, the intracellular calcium chelator BAPTA-AM and the Ca^2+^ ionophore A23187 are shown. **a**, Viability. **b**, Proliferation. **c**, Switching. Values represent mean ± s.e.m. of three independent experiments performed in triplicate. **p*<0.05; ***p*<0.01; ****p*<0.001. *****p*<0.0001; ns, not significant.
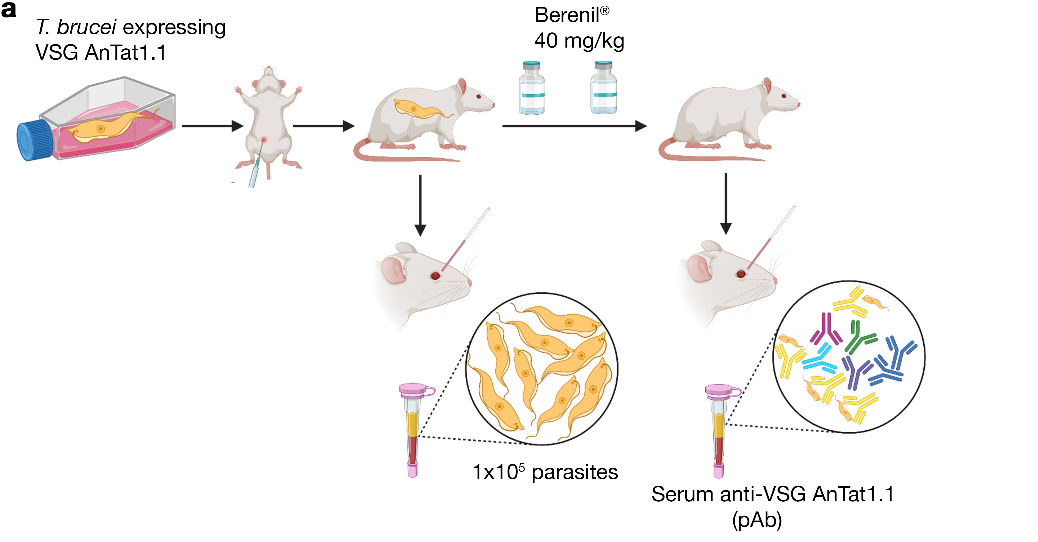

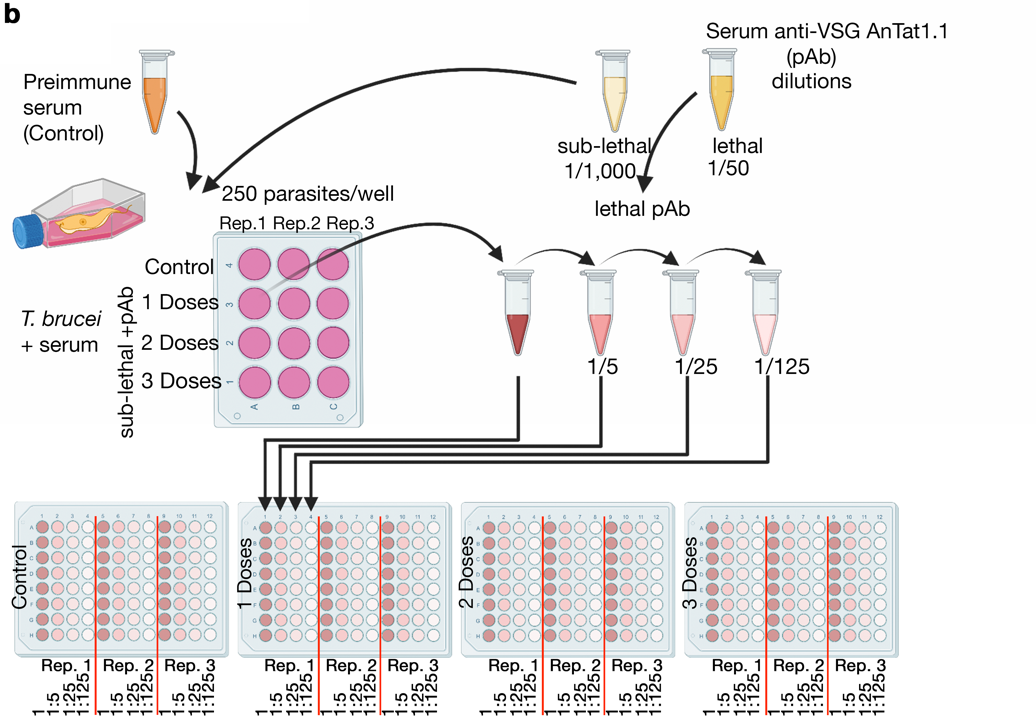

**Extended Data Figure 6| Schematic representation of *in vitro* induction of switching in *T. brucei*.** **a**, Cloned parasites expressing VSG AnTat1.1 were grown in culture flasks and used to infect mice from which pre-immune serum was previously collected (control serum). After 2 dpi, blood was taken and the presence of less than 1x10^5^ parasites/ml of serum was verified to avoid the appearance of switchers. Infections were immediately treated with a dose of Berenil^®^, which was repeated 24 h later for complete cure of the animals. At 15 dpi, total blood was collected from the animals and the immune serum (pAb) was isolated and titrated to determine sub-lethal and lethal dilutions on *T. brucei* parasites expressing VSG AnTat1.1. **b**, Approximately 250 parasites/ml were distributed in 12-well plates in triplicate and confronted with daily sub-lethal doses (1/1,000) of immune serum or pre-immune serum (1/50) as control. After 3 days, parasites from each well were counted to control growth rates and then serially diluted and incubated with a lethal dilution of the immune serum (1/50) to kill non-switchers. After 6 days, clones from randomly selected wells were isolated to determine the expressed VSG by RT-PCR and cDNA sequencing. Results are shown in Fig. 4g.

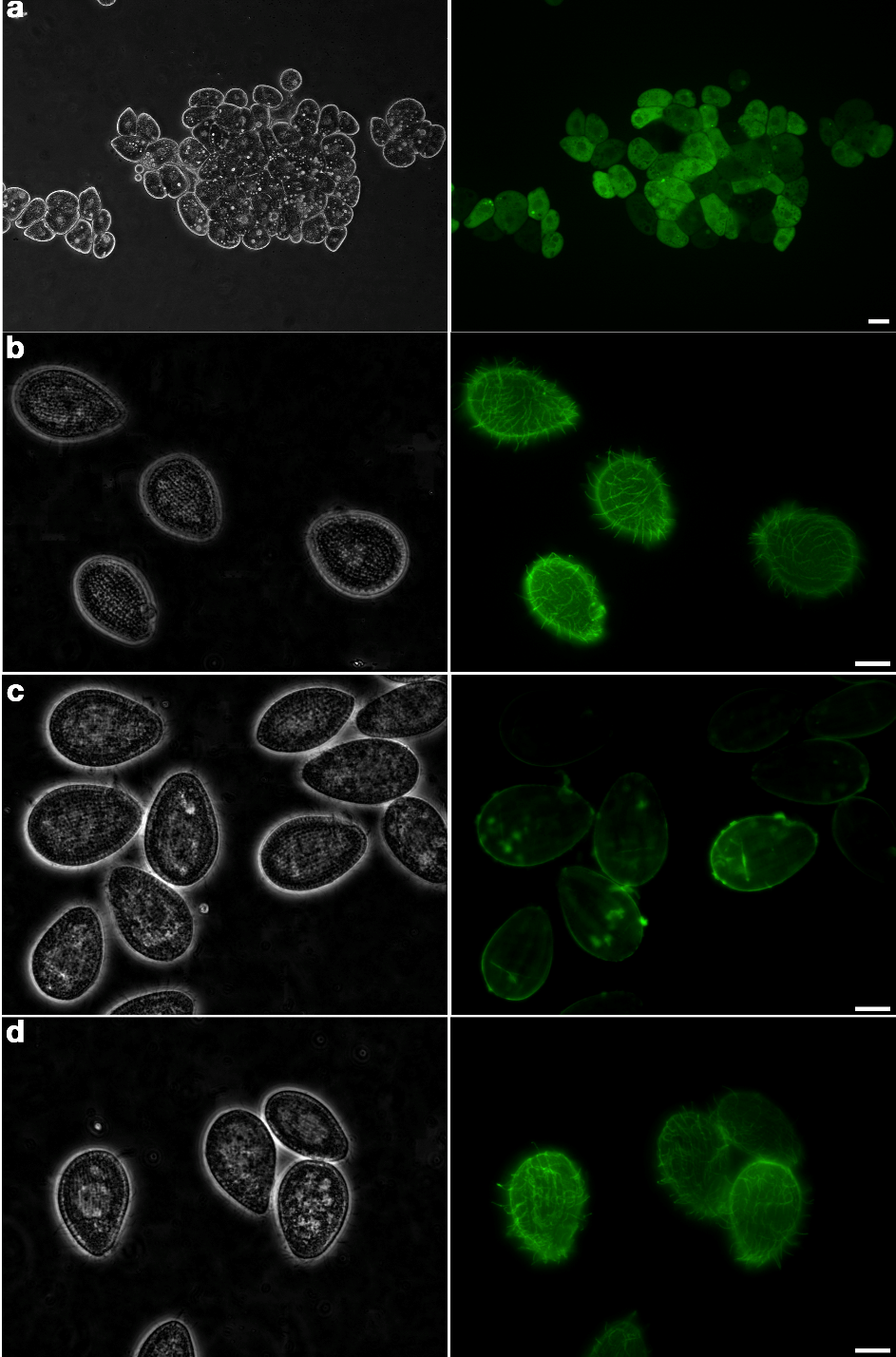

**Extended Data Figure 7| Effect of antiserum directed to *T. thermophila* surface antigens.** *T. thermophila* cells treated with homologous antiserum (1/1,000) and analysed for cell viability using FDA/PI staining and IFA to detect the expression and localization of the original surface antigen. In either case, the images on the left correspond to phase contrast microscopy and images on the right to either FDA/PI staining (A) or immunofluorescence labelling with a secondary antibody (B). **a**, *T. thermophila* cells form large aggregates of live cells after 24 h in contact with a polyclonal serum directed to their immobilization-antigens (green cytoplasm labelled with FDA). **b**, Control of untreated cells. The entire surface of the cells, including cilia, is labelled. **c**, Cells treated for 24 h in the presence of the antiserum where labelling of the cilia has highly diminished. **d**, Cells treated for 24 h with antiserum and in the absence of the antibodies for 4 additional days. Scale bars, **a**, 50 µm; **b**, 10 µm.

**Supplementary Movie S1| Anti-VSP antibodies induce rapid detachment and agglutination of trophozoites without killing the parasites.** Time-lapse observation of *Giardia* trophozoites expressing VSP417 during incubation with 100 µM of mAb 7C2. It can be seen how trophozoites quickly detach from the wall of the glass culture tubes and form large aggregates containing live cells. These large clumps of motile trophozoites remain unmodified for up to 3 days. (See separate .avi file)
